# Refactored intracellular domains enhance CAR expression and function

**DOI:** 10.64898/2026.10.28.762481

**Authors:** Liam Bai, Aakriti Kumar, Hetvi Desai, Jiahe Li, Dinesh de Alwis, Daria Hazuda, Gevorg Grigoryan, Wujie Wang, Jason Yi

## Abstract

Chimeric antigen receptor (CAR) intracellular domains (ICDs) couple antigen recognition to T-cell activation. Here we worked with AI agents to refactor CAR ICDs, proposing designs and refining them as measurements accumulated over three rounds. This approach yielded `krgr–ζ1–40`, a 45-amino-acid ICD less than a third the length of the clinical `4-1BB–CD3ζ` (`BBζ`) ICD. Compared with `BBζ`, `krgr–ζ1–40` increased CAR surface expression and antigen-driven IFN-γ release, without detectable antigen-independent IFN-γ release. `krgr–ζ1–40` extended CAR expression and target-cell killing after transient mRNA delivery, supported the expression of four CARs from a shared mRNA dose, and enhanced B-cell depletion after lentiviral and lipid-nanoparticle delivery. These results show that iterative agent-guided design with experimental feedback can yield compact ICDs that enhance CAR expression and sustain CAR T-cell function.

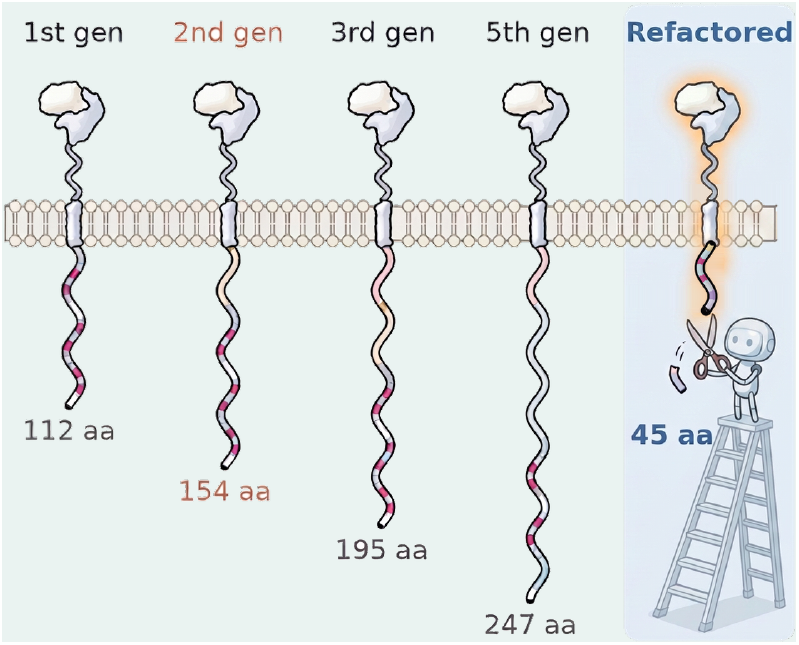

## Introduction

Chimeric antigen receptors (CARs) redirect T cells to recognize cell-surface antigens independently of major histocompatibility complexes (MHCs). The concept originated with chimeric T-cell receptors whose variable regions were replaced by those of an antibody, which conferred antibody-type specificity on T cells [1]. In parallel, Irving and Weiss showed that the cytoplasmic domain of the T-cell receptor (TCR) ζ chain is sufficient to couple a chimeric receptor to TCR signal transduction pathways [2]. Combining these ideas, Eshhar and colleagues fused a single-chain antibody fragment to the ζ or γ chain, producing single-chain chimeric receptors that triggered cytokine secretion and target-cell lysis upon antigen binding [3]. Adding a costimulatory domain from CD28 [4,5] or 4-1BB [6–8] to CD3ζ produced the second-generation CARs used in approved therapies, and the two differ in signaling, metabolism and persistence [9,10].

Engineering these receptors requires balancing antigen-induced activity with behavior in the absence of antigen. Antigen-independent (tonic) CAR signaling can promote exhaustion, and 4-1BB and CD28 costimulation have different effects on exhaustion in tonic-signaling models [11]. Activation strength also depends on the number of immunoreceptor tyrosine-based activation motifs (ITAMs) in CD3ζ. Mutating two of the three CD3ζ ITAMs lowers the activation potential of a CD28-based CAR, which preserves memory differentiation and limits exhaustion [12]. Conversely, additional ITAMs increase T-cell proliferation [13,14] and the efficiency of receptor triggering, likely by bringing multiple ZAP-70 molecules into proximity [15], and the larger number of ITAMs in the TCR–CD3 complex is thought to contribute to the greater antigen sensitivity of the TCR compared with CARs [15,16]. These studies support the prevailing view that more ITAMs produce stronger activation, so that removing ITAMs trades acute activity for persistence. They also suggest that the composition of the CAR intracellular domain (ICD) can be tuned beyond the two clinical designs, `CD28–CD3ζ` and `4-1BB–CD3ζ`.

Efforts to expand this signaling vocabulary have screened libraries of signaling domains for T-cell function, from randomly assembled domains to pooled and combinatorial designs [17–24]. Pooled screens have tested dozens of intracellular domains from diverse immune receptors in primary T cells [19–21], shuffled costimulatory domains with single-cell readouts [22–24] and recombined short signaling motifs to decode their effects on CAR T-cell phenotype [17]. Others have built CARs from CD3 subunits beyond CD3ζ [25,26], from the ITAM adaptor DAP12 [27] and from proximal signaling molecules [28,29]. Together, these studies show that the signaling vocabulary of CARs extends beyond CD28 and 4-1BB. However, they explored which domains to combine, exchange or reorder rather than how much of the ICD could be removed. The closest test, truncating CD3ζ to a single membrane-proximal ITAM in a CD28-based CAR, preserved acute effector function and improved tumor control, but was designed to examine ITAM position and dosage and retained the full CD28 domain [12]. Because shortening the signaling domain is otherwise expected to reduce function, how far the ICD as a whole can be compacted, and whether compaction can improve function, remain unknown.

Refactoring in software reorganizes internal structure while preserving observable behavior [30]. In biology, refactoring has simplified systems shaped by natural selection, from the overlapping genetic elements of bacteriophage T7 to G protein-coupled receptor signaling in yeast, so that they can be engineered rationally [31,32]. Applied to CAR ICDs, refactoring would reorganize signaling motifs into compact domains that preserve the core function of coupling antigen recognition to T-cell activation. This requires choosing which binding partners to recruit, how to combine and place their motifs, and which sequences to remove. Unlike the folded proteins of structure-based design [33–35], signaling domains such as the CD3ζ cytoplasmic domain are largely disordered and act through short motifs that recruit partners inside cells [17,36,37]. The functional consequences of these design choices must be established experimentally. AI agents that combine literature search, data analysis and hypothesis generation to propose and revise experiments [38,39] offer a way to reason over the design space as measurements accumulate.

A compact ICD would also remove most of this disordered sequence, which two mechanistic models implicate in receptor signaling, with opposite predictions for a compact ICD. In the first, signaling components assemble into membrane condensates that promote signal transduction [40,41]. In T cells, these condensates have been attributed mainly to multivalent crosslinking of LAT [40,42], as in other multivalent signaling systems [43]. However, TCR– ZAP-70 and LAT occupy distinct domains within microclusters, with the TCR–ZAP-70 domain assembling first [44], and LAT condenses only after a delay following TCR engagement [44,45], so LAT crosslinking does not explain how the receptor itself clusters. In the broader condensate field, phase separation is often driven by intrinsically disordered regions (IDRs) themselves, in which interacting stickers, such as aromatic and charged residues, are separated by solvating spacers [46–48], and in CARs, an engineered CD3ε motif that condenses through cation–π interactions improved antigen sensitivity and persistence [49]. If the receptor’s disordered tails drive condensation and condensation drives activation, removing disordered sequence should impair activation. In the second model, basic-rich stretches (BRS) of the CD3ε and CD3ζ tails bind acidic phospholipids of the inner leaflet [50–57], inserting ITAM tyrosines into the bilayer and restraining their phosphorylation until antigen engagement releases them [50,51,55], although disrupting the CD3ζ BRS has also been found to weaken TCR signaling or synapse accumulation [52,54]. Removing the BRS should instead release tonic signaling. A compact ICD therefore risks either loss of activation or loss of antigen dependence, and changing disorder, basic residues or motif spacing can test these competing expectations.

Compactness may also change how many receptors reach the cell surface. CAR function depends on both receptor and antigen density [58–61], so a compact ICD that raises surface expression could improve function even without greater signaling per receptor. Transient mRNA delivery, which can generate functional CAR T cells [62,63], provides a setting in which this relationship can be followed as receptors are lost from the cell surface. Expression from limited mRNA also matters for in vivo CAR T-cell generation, in which targeted lipid nanoparticles carrying CAR mRNA [64–66] or targeted lentiviral vectors [67] modify T cells inside the body. With lipid nanoparticles, an ICD that yields more receptor from less mRNA could lower the dose needed for function.

Here we addressed these questions by refactoring CAR ICDs through successive rounds of design, in which scientists and AI agents proposed changes to motif composition and sequence context and the resulting measurements guided the next designs. We asked how far the ICD could be compacted and whether removing its disordered sequence would cost activation or antigen dependence, as the condensate and lipid-binding models predict. We also asked whether a compact ICD that yields more surface receptor would improve function after mRNA and lipid-nanoparticle delivery, and whether experimental feedback could steer design ideas across rounds. Successive rounds produced shorter ICDs with higher expression and function, without the losses that either model predicts. Earlier-round measurements also shifted large language model (LLM) predictions toward the design ideas that worked, although measurement still revealed gains the models missed. These results establish iterative design by scientists and AI agents as a route to compact CAR ICDs with higher expression and more durable function.

## Results

### Agent-guided design and screening identify brs–ζ1–40, a compact CAR ICD

CAR intracellular domains are typically assembled from intact cytoplasmic domains of CD3ζ and a costimulatory receptor such as 4-1BB or CD28, yet T cells use a much larger vocabulary of signaling modules [17,19,20,22,25,68– 71]. To make this vocabulary explicit and to frame the design of compact ICDs, we compiled published evidence into a signaling map (Fig. 1a). The map links 34 ICD modules, drawn from CD3 chains and other ITAM receptors, co-receptors, the CD28 family, LAT, the TNF-receptor family and cytokine receptors, through binding partners and relays to signaling pathways and reported T-cell phenotypes. It draws on 150 interactions from 253 primary studies, each graded by the strength of its evidence [68,72–75]. Our iterative exploration across rounds 1–3 covered most modules on the map (Fig. 1a).

**Figure 1.**
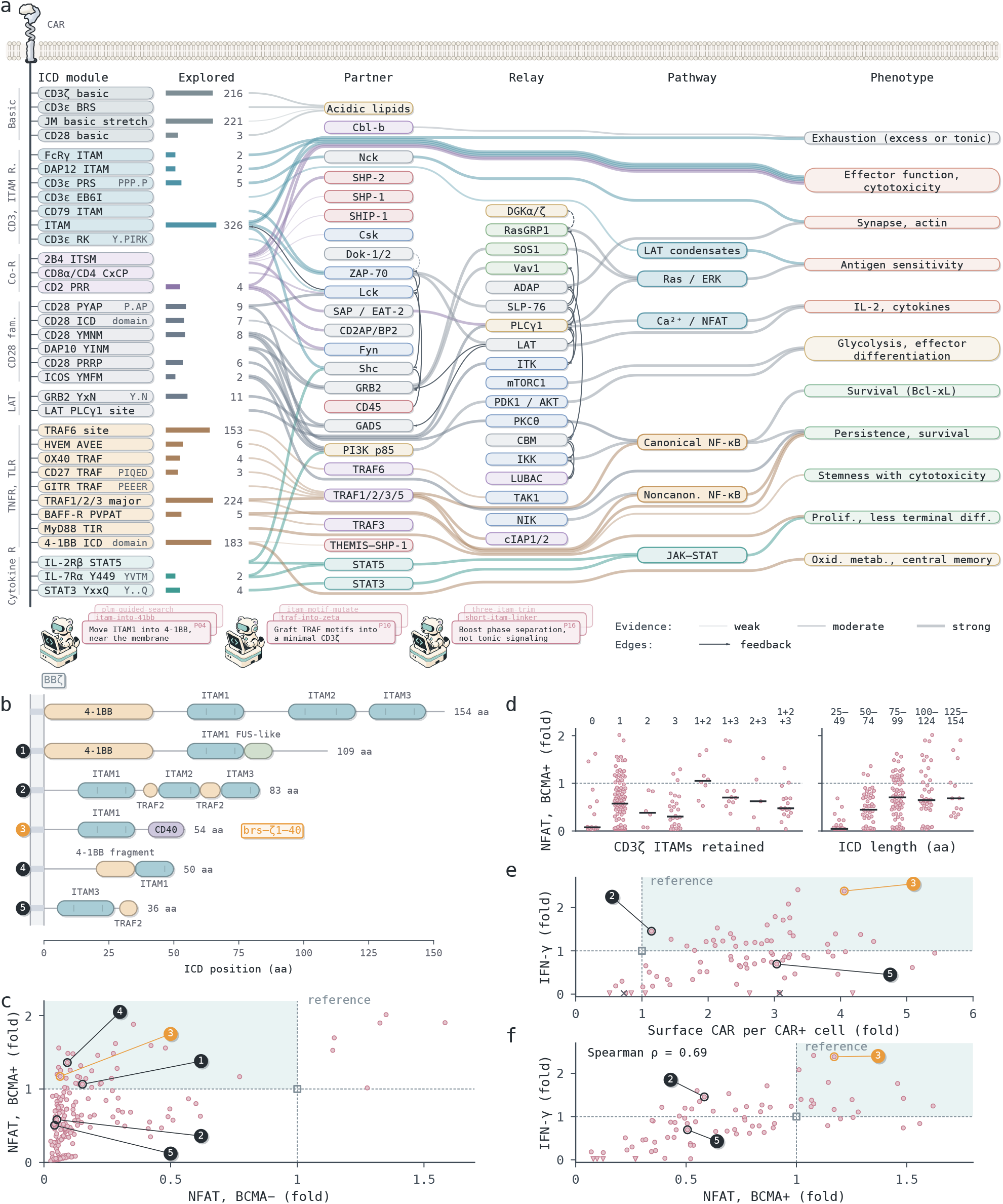
Agent-guided design and screening of compact ICDs. **a**, Signaling map of ICD modules. Designs compose sequence motifs that recruit partner proteins, which relay to signaling pathways. Lines show the strong activating interactions, plus the best-supported interaction of any otherwise unconnected node. Explored, number of the 339 ICDs tested in rounds 1–3 that carry each module’s sequence motif (bar length on a log scale); blank, none. **b**, Representative designs proposed by scientists and agents, drawn to residue scale. **c**, Many compact designs show higher on-target signaling and lower background signaling than `BBζ` (`4-1BB–CD3ζ`). Y axis shows on-target activation in BCMA-positive cells; X axis shows background activation in BCMA-negative cells; expressed as fold over reference (open square). Shading marks higher on-target and lower background signal than `BBζ`. **d**, A single ITAM can support strong signaling. Jurkat NFAT signal expressed as fold of `BBζ`, grouped by native CD3ζ ITAMs retained and by ICD length; bars, medians. **e**, Designs re-tested in primary T cells: most showed higher surface expression than `BBζ`, and some released more IFN-γ. Surface CAR density and antigen-driven IFN-γ, expressed as fold of `BBζ`; ×, ITAM Y→F and no-mRNA negative controls; shading marks higher surface CAR and IFN-γ than `BBζ`. **f**, Correlation between Jurkat NFAT and primary T-cell IFN-γ for the 78 re-tested ICDs. Each axis is relative to its assay’s `BBζ` reference (open square; dashed lines); shading marks higher NFAT and IFN-γ than `BBζ`. Spearman ρ = 0.69 across all 78 designs.

The map summarizes established rules of CAR signaling. CD3ζ ITAM number and identity calibrate activation strength, ligand sensitivity and T-cell fate [12,15,76]. 4-1BB signals through TRAF1–3 and noncanonical NF-κB [77,78] and recruits THEMIS–SHP-1, which attenuates Lck-dependent priming [79]. CD28 and 4-1BB differ in the kinetics, magnitude and metabolic consequences of signaling [9,10], and point mutations in CD28 motifs change persistence and function [80,81]. Membrane-proximal basic residues bind acidic inner-leaflet lipids [51,52]; CD3ε motifs recruit Lck, PI3K and Csk [82–84]; and a JAK–STAT module can add a cytokine signal [85]. ICD size, rather than costimulatory identity, determines constitutive CD3ζ phosphorylation [86], and pooled and combinatorial libraries show that non-canonical domains and module combinations can change CAR T-cell phenotype [17,19,20,22]. Grading the evidence behind each rule family by experimental system showed that most support comes from T cells in vitro and from mouse models, whereas clinical evidence is limited to comparisons of CD28 and 4-1BB costimulation.

We built an agent system that combines LLM reasoning with literature search and computational tools. Scientists specified design objectives and reference sequences, and tool-using agents searched the literature and proposed sequence changes. Scientists then reviewed the proposed designs, steered the agents and compared candidate sequences. After several cycles of review we selected 186 candidates for the round 1 library screen. Representative designs ranged from 36 to 109 amino acids (aa), compared with 154 for the second-generation `4-1BB–CD3ζ` reference ICD (`BBζ`; Fig. 1b).

As an initial exploration, we screened the 186 designs as CARs with the D-domain BCMA binder (ddBCMA) [87] in Jurkat NFAT reporter cells against BCMA-positive and -negative target cells. Of the 186 designs, 38 matched or exceeded the BCMA-driven signal of `BBζ`, and 32 of these also gave lower signal with BCMA-negative targets (Fig. 1c). Grouping the designs by native ITAM count and length, we found that 25 of the 130 designs retaining only one ITAM supported antigen-driven signaling at or above the level of `BBζ` (Fig. 1d).

`brs–ζ1–40` is a 54-aa ICD joining the membrane-proximal basic clusters and first ITAM of CD3ζ to a CD40 segment required for TRAF6 binding in native CD40 [88,89]. CD40-derived signaling domains have previously enhanced CAR T-cell proliferation, survival and persistence [23,90,91]. However, a 19-residue CD40 motif spanning the same TRAF6-binding site was among the most inhibitory motifs for CAR T-cell proliferation in a pooled screen of CARs with full-length CD3ζ [17], so whether this segment can act as a costimulatory domain within this scaffold was unclear. Unexpectedly, `brs–ζ1–40` matched the NFAT signal of `BBζ` with BCMA-positive targets, whereas with BCMA-negative targets its signal was about one-sixteenth that of `BBζ` (Fig. 1c).

To test whether the designs transferred to another binder and to primary T cells, we re-screened 78 designs, selected by ICD length and NFAT activity, as bb2121 BCMA scFv CARs [92] in primary T cells (Fig. 1e). Strikingly, nearly all compact designs increased surface CAR: 73 of the 78 exceeded `BBζ` (median, 2.7-fold), and 30 also released more antigen-driven IFN-γ. In this screen, `brs–ζ1–40` displayed 4.1-fold the surface CAR of `BBζ` and the second-highest IFN-γ of the 78 designs. Without target cells, its IFN-γ remained below the detection limit. Jurkat NFAT and primary T-cell IFN-γ ranked the re-screened designs similarly (Spearman ρ = 0.69; Fig. 1f), even though the two screens differed in binder, target cells and measured functional response. Together, these results identify `brs–ζ1–40` as a compact ICD that is highly expressed and antigen responsive in primary T cells.

### brs–ζ1–40 increases CAR expression across BCMA binders and retains antigen-dependent function

We selected ten compact ICDs, including `brs–ζ1–40`, to test in primary T cells across a range of effector-to-target (E:T) ratios (Fig. 2a–c). All ten compact ICDs displayed more surface CAR than `BBζ`, from 2.3- to 17-fold (Fig. 2a).

**Figure 2.**
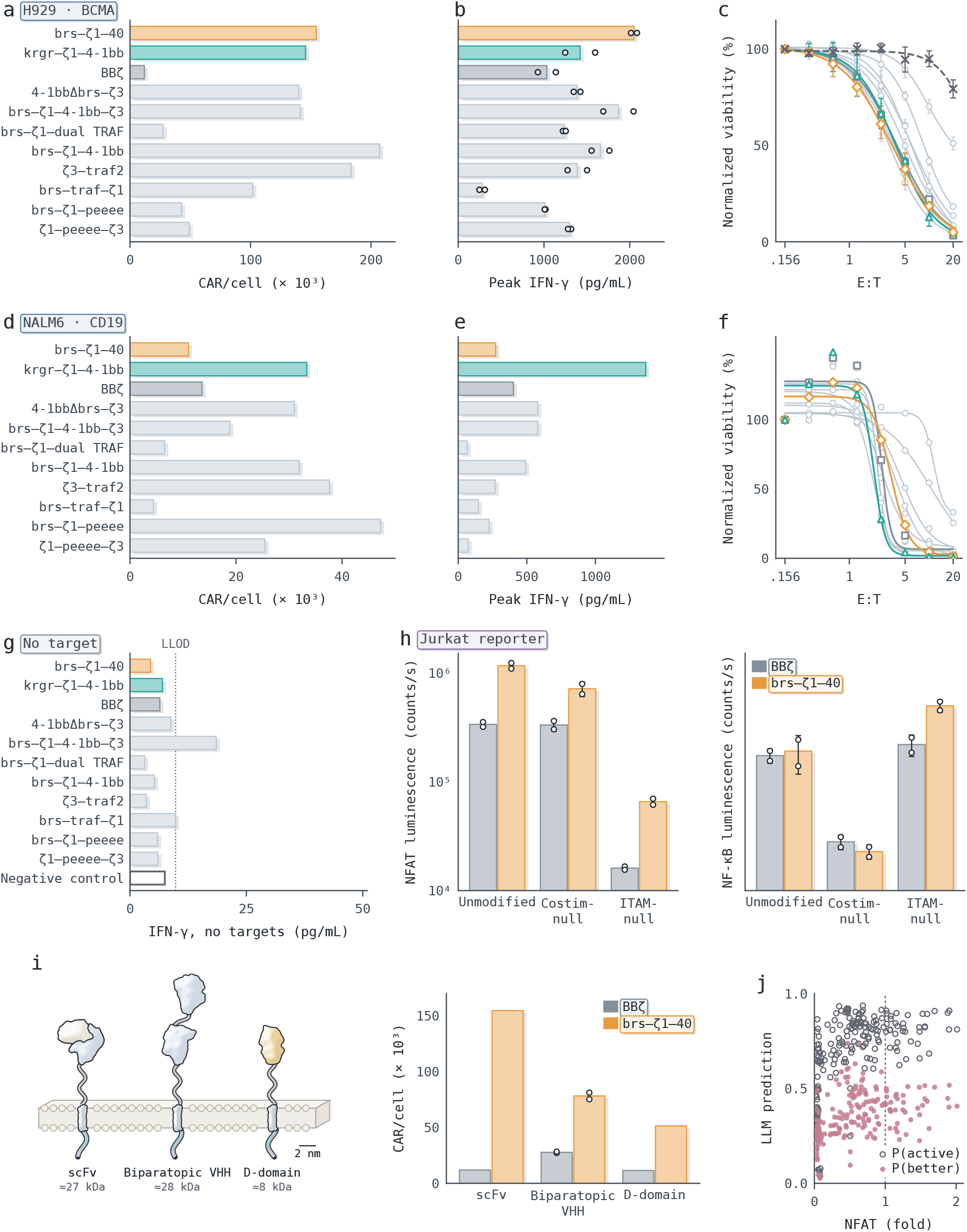
Compact ICD expression, killing and signaling across CAR binders. **a–c**, Surface CAR density (a), peak IFN-γ across eight E:T ratios (b) and killing of H929 myeloma cells (c) by primary T cells expressing bb2121 BCMA CARs. **d–f**, The same ICDs in FMC63 CD19 CARs: surface CAR density (d), peak IFN-γ across eight E:T ratios (e) and killing of NALM6 leukemia cells (f). Rows in b and e follow a and d. Killing is shown as residual target-cell luminescence normalized to the lowest E:T ratio for each construct and replicate; lower values indicate more killing. **g**, Peak IFN-γ from BCMA CAR T cells without targets across eight effector inputs; dotted line (LLOD), 9.8 pg/mL lower limit of detection. **h**, Jurkat NFAT and NF-κB responses to costimulation- or ITAM-disrupting mutations in `brs–ζ1–40` and `BBζ` (`4-1BB– CD3ζ`). In `brs–ζ1–40`, E42A/E45A substitutions alter the CD40-derived TRAF6-binding segment (QEPQE→QAPQA), and Y21F/Y32F substitutions replace both ITAM1 tyrosines. The corresponding `BBζ` mutants disrupt 4-1BB costimulation or replace all six tyrosines across the three CD3ζ ITAMs with phenylalanine. **i**, Three binder formats, and `brs–ζ1–40` vs. `BBζ` expression across them. **j**, Zero-shot LLM predictions for the round 1 library against NFAT relative to `BBζ` (Fig. 1c); mean of five models.

We next asked whether this higher expression was accompanied by cytokine release and target-cell killing. CAR T cells expressing the compact designs were co-cultured with BCMA-positive H929 myeloma cells across an E:T titration. `brs–ζ1–40` released the most IFN-γ in the panel, 2.0-fold the peak of `BBζ` (Figs. 2b and S1e), and ranked highest on both replicate plates (Fig. S1d). Several designs killed H929 cells as effectively as `BBζ`, whereas the design with a TRAF motif inserted before ITAM1 (`brs–traf–ζ1`) killed only about half of the H929 cells even at the highest E:T (Fig. 2c). `brs–ζ1–40` and `BBζ` killed H929 cells with similar potency. These results show that `brs–ζ1–40` releases more IFN-γ than `BBζ` while killing BCMA-positive target cells as effectively.

To determine whether these properties depend on the binder, we tested the same eleven ICDs with the FMC63 anti-CD19 scFv [93] against NALM6 leukemia cells (Fig. 2d–f). Seven of the ten compact designs again displayed more surface CAR than `BBζ`, by up to 3.5-fold (Fig. 2d). In contrast, `brs–ζ1–40` displayed slightly less CAR than `BBζ` with FMC63, whereas `krgr–ζ1–4-1bb` displayed 2.5-fold more. `brs–ζ1–40` released less NALM6-induced IFN-γ than `BBζ`, whereas `krgr–ζ1–4-1bb` released the most in the panel, 3.4-fold `BBζ` (Figs. 2e and S1f). `brs–ζ1– 40` and `BBζ` killed NALM6 cells with similar potency (Fig. 2f), and `krgr–ζ1–4-1bb` was among the most potent. Thus, the expression and cytokine advantage of `brs–ζ1–40` seen with bb2121 did not extend to FMC63, whereas `krgr–ζ1–4-1bb` was highly expressed and active with both binders.

Because tonic signaling can promote exhaustion [11] and may rise with receptor density [59,94], we next measured IFN-γ released by the BCMA CAR T cells in the absence of target cells (Fig. 2g). Across effector inputs, every construct except `brs–ζ1–4-1bb–ζ3` remained below the 9.8 pg/mL detection limit, including `brs–ζ1–40` despite its 13-fold higher receptor density. These results indicate that the high surface density of `brs–ζ1–40` does not lead to detectable tonic IFN-γ release.

To determine which segments of `brs–ζ1–40` drive its signaling, we compared signaling mutants of `brs–ζ1–40` and `BBζ` in Jurkat NFAT and NF-κB reporter cells stimulated with BCMA-positive target cells (Fig. 2h). Costimulation-null variants carried alanine substitutions in the CD40-derived QEPQE motif of `brs–ζ1–40` or the TRAF-binding motif of 4-1BB, and ITAM-null variants carried Y→F substitutions in every ITAM. As expected, the costimulation-null variant of `BBζ` kept its NFAT signal but lost more than 80% of its NF-κB signal, whereas the ITAM-null variant lost about 95% of its NFAT signal. `brs–ζ1–40` followed a similar pattern: its costimulation-null variant lost more than 80% of NF-κB signal but, unlike that of `BBζ`, also about 40% of NFAT signal, and its ITAM-null variant lost about 95% of NFAT signal. In this experiment, unmodified `brs–ζ1–40` gave 3.5-fold higher NFAT and similar NF-κB signal compared with `BBζ`, more than the matched NFAT signal of the library screen (Fig. 1c). These results show that, as in `BBζ`, NFAT signaling by `brs–ζ1–40` depends on its ITAM tyrosines and NF-κB signaling on its CD40-derived QEPQE motif.

We then asked whether the expression advantage of `brs–ζ1–40` extends to other BCMA binder formats represented among clinical CAR T cells (Fig. 2i). With the biparatopic VHH binder of ciltacabtagene autoleucel, `brs–ζ1–40` displayed 2.8-fold more CAR per cell than `BBζ` at similar CAR-positive fractions. With ddBCMA, it displayed 4.5-fold more, compared with 13-fold with the bb2121 scFv (Fig. 2a). Together, these results show that `brs–ζ1– 40` increases surface CAR expression across scFv, VHH and miniprotein BCMA binders while retaining antigen-dependent killing and signaling that depends on its ITAM and its CD40-derived QEPQE motif.

We also asked whether such designs could have been anticipated from sequence alone. Given each library sequence and a description of the assay, five general-purpose LLMs predicted NFAT activity poorly. Their predicted probability that a design would exceed `BBζ` in NFAT stayed below 0.75 and hardly separated the 38 designs that did from the rest (mean 0.40 versus 0.34; Fig. 2j).

### Experiment-informed agent redesign yields krgr–ζ1–40, a 45-aa ICD with higher expression and IFN-γ

To determine which features of the round 1 hits could be combined in a shorter ICD, we worked with Claude Code to design 15 round 2 ICDs from four hits characterized in primary T cells (Fig. 3a). The designs tested six groups of changes: ablations, costimulatory motif swaps, ITAM swaps, two TRAF-binding motifs, KRGR anchor transfers and minimal domains. Ablations tested whether individual segments were required, while motif swaps and anchor transfers tested whether features of different hits could be combined; `krgr–ζ1–40` was an anchor transfer. `brs–ζ1–40` had released the most IFN-γ of the four hits, and earlier cross-binder cytokine measurements motivated transferring the KRGR anchor from `krgr–ζ1–4-1bb` (Fig. 3a). The Jurkat screen had also shown that KRGR was required for the activity of `krgr–ζ1–4-1bb` (Fig. S1b). These observations led us to replace the first 13 residues of the basic-rich segment of `brs–ζ1–40` with the four-residue KRGR anchor while retaining its ITAM and CD40-derived segment (Fig. 3c). The resulting `krgr–ζ1–40` was 45 aa long, shorter than either parent (54 and 56 aa) and less than a third the length of `BBζ`.

**Figure 3.**
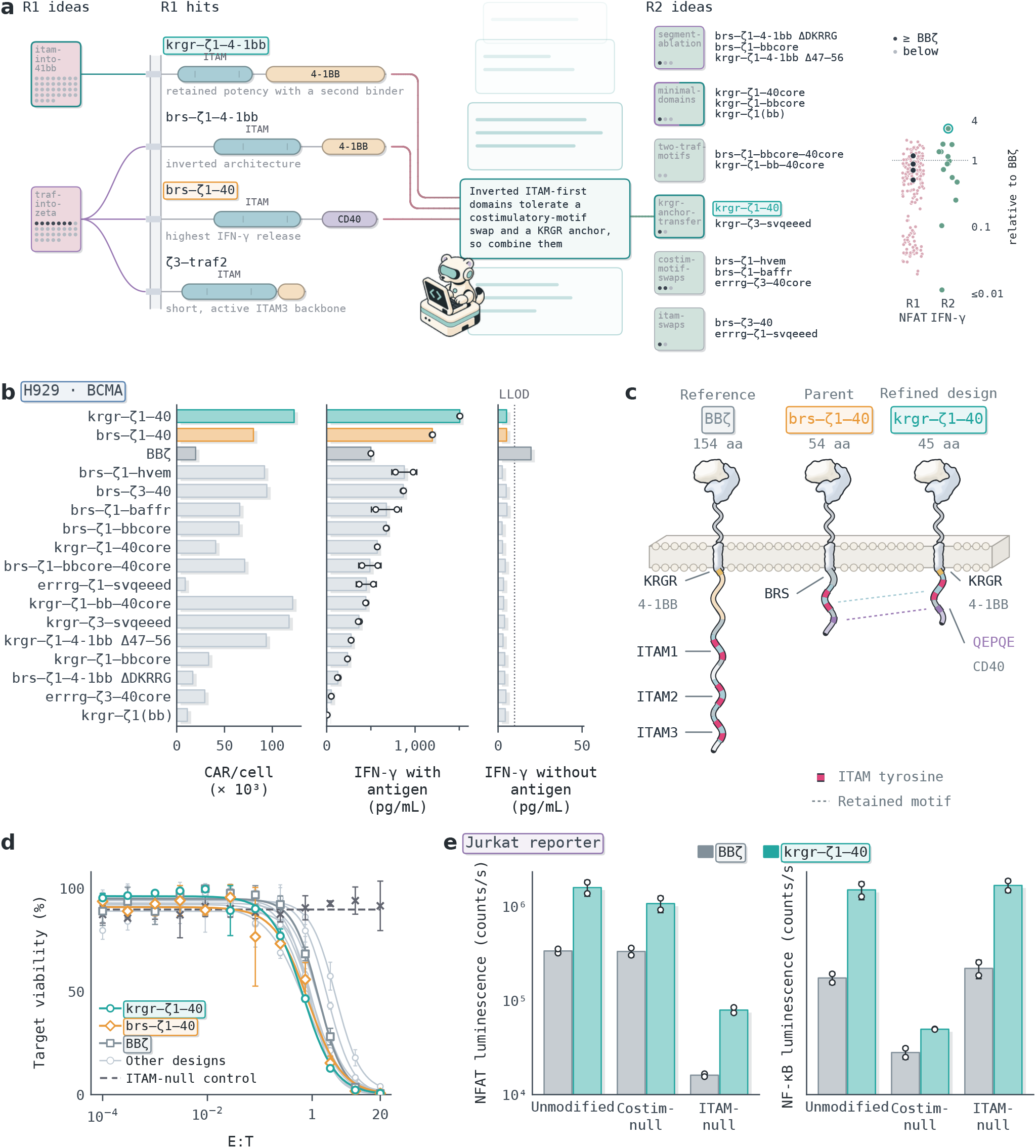
krgr–ζ1–40 refinement and functional characterization. **a**, Evolution of design ideas from round 1 to round 2. Four round 1 hits were the parents of the 15 round 2 designs, which recombine sequence features from these hits. Center, a simplified reasoning step by which the agent drew on three round 1 hits to combine the KRGR anchor of `krgr–ζ1–4-1bb` with the ITAM and CD40-derived segment of `brs–ζ1–40`. Right, round 1 and round 2 activation readouts relative to `BBζ` (`4-1BB–CD3ζ`), in Jurkat and primary T cells respectively. **b**, Round 2 results in H929 cells with the bb2121 binder: CAR density, IFN-γ and IFN-γ without antigen. **c**, Reference `BBζ` (154 aa), parent `brs–ζ1–40` (54 aa) and refined `krgr–ζ1–40` (45 aa). **d**, Killing of H929 myeloma cells by round 2 designs, including `krgr–ζ1–40`, `brs–ζ1–40` and `BBζ` across E:T ratios. **e**, Jurkat NFAT and NF-κB responses to costimulation- or ITAM-disrupting mutations in `krgr–ζ1–40`. In `krgr– ζ1–40`, E33A/E36A substitutions alter the CD40-derived TRAF6-binding segment (QEPQE→QAPQA), and Y12F/Y23F substitutions replace both ITAM1 tyrosines.

To determine whether the KRGR anchor preserved the properties of `brs–ζ1–40`, we tested the 15 round 2 designs alongside `brs–ζ1–40` and `BBζ` in a primary T-cell screen (Fig. 3b). Twelve of the 15 round 2 designs displayed more surface CAR than `BBζ`, and six released more antigen-driven IFN-γ. `krgr–ζ1–40` improved on the surface expression of `brs–ζ1–40` by 1.5-fold and `BBζ` by 6.2-fold (Fig. 3b, left). Its IFN-γ was the highest of the 15 round 2 designs (Fig. 3b, middle). Without target cells, all round 2 designs and `brs–ζ1–40` remained below the detection limit (Fig. 3b, right). Thus, replacing the 13-residue basic-rich segment of `brs–ζ1–40` with the four-residue KRGR anchor increased both expression and antigen-driven IFN-γ without increasing tonic IFN-γ.

To compare killing, we titrated `krgr–ζ1–40`, `brs–ζ1–40` and `BBζ` CAR T cells against H929 cells across a range of E:T ratios (Fig. 3d). Killing curves for `krgr–ζ1–40` and `brs–ζ1–40` were left-shifted relative to that of `BBζ`, at 2.0- and 1.6-fold lower IC50 E:T, respectively, so fewer CAR T cells were needed per target cell for the same killing.

To determine whether `krgr–ζ1–40` signals through the same segments as `brs–ζ1–40`, we tested the signaling mutants of Fig. 2h (Fig. 3e). Unmodified `krgr–ζ1–40` gave 4.7-fold higher NFAT and 8.6-fold higher NF-κB signal than `BBζ`, in proportion to its higher surface CAR (Fig. S2). In contrast, `brs–ζ1–40` gave about a quarter of the NF-κB signal per surface CAR of `BBζ`, which may reflect suboptimal costimulatory signaling (Fig. S2). As with `brs–ζ1–40`, NFAT signaling depended on the ITAM tyrosines, with about 95% lost in the ITAM-null variant, and NF-κB signaling on the CD40-derived motif, with more than 95% lost in the costimulation-null variant, compared with about 85% for `BBζ` (Fig. 3e). Together, these results show that the KRGR anchor preserves the ITAM- and CD40-dependent signaling of `brs–ζ1–40` while further increasing expression and antigen-driven IFN-γ. We therefore carried `krgr–ζ1–40` forward as the lead compact ICD.

### krgr–ζ1–40 extends CAR expression and function across mRNA, lentiviral and lipid-nanoparticle delivery

We next measured how `krgr–ζ1–40` expression scales with the amount of mRNA delivered and how long expression and function last after transient delivery (Fig. 4a–e). Surface CAR density increased with the mRNA dose for all three ICDs, but `krgr–ζ1–40` gave the highest density at every dose, 8.4-fold that of `BBζ` and 2.4-fold that of `brs–ζ1–40` at the highest dose (Figs. 4a and S3). `krgr–ζ1–40` matched the density that `BBζ` reached at the highest dose with one-eighth of the mRNA.

**Figure 4.**
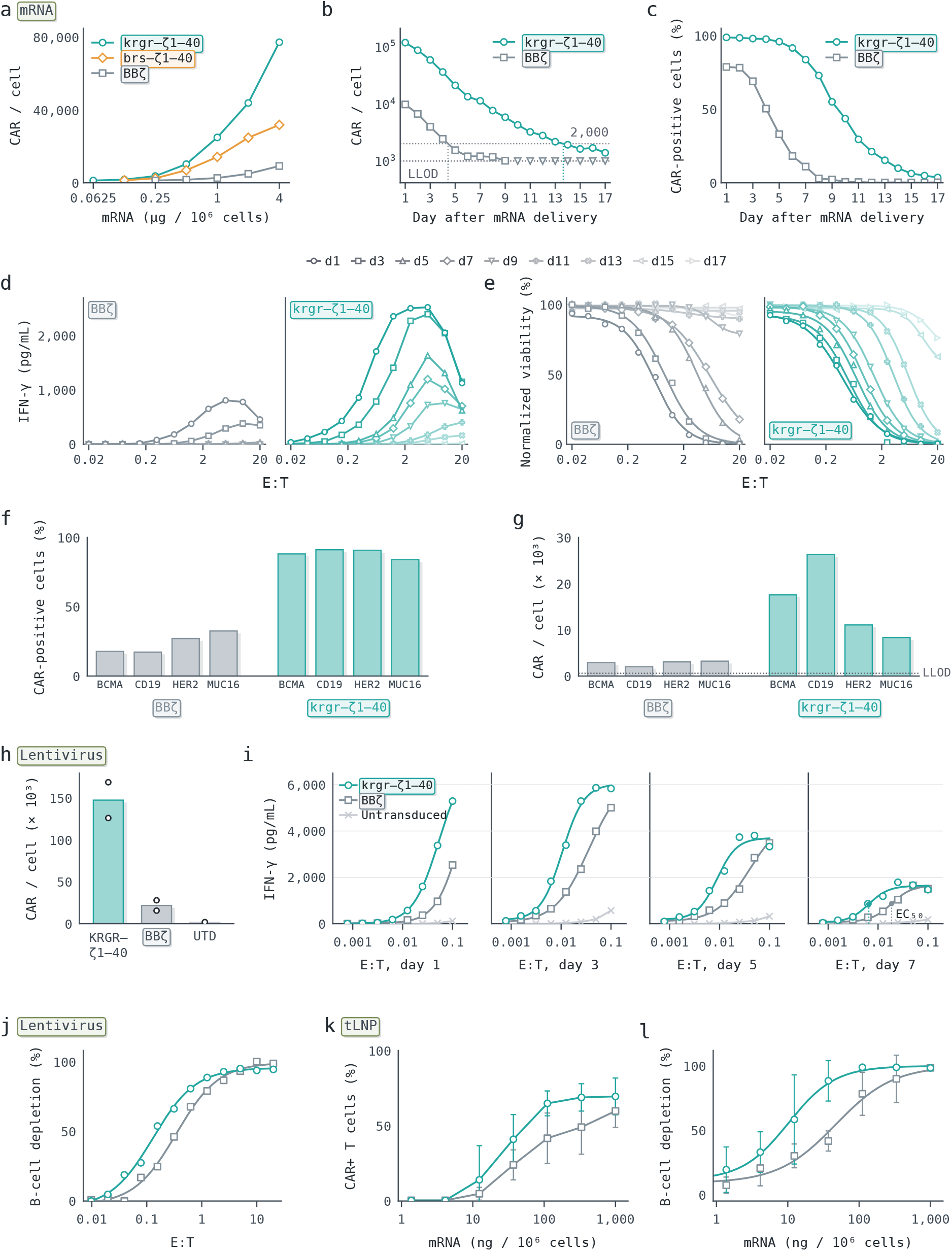
krgr–ζ1–40 expression, persistence and function after mRNA electroporation, lentiviral and tLNP delivery. **a**, Surface CAR density 24 h after electroporation with increasing CAR mRNA doses. **b**, Surface CAR density over 17 days after mRNA delivery. **c**, CAR-positive cells over the same period. **d**, IFN-γ across E:T ratios on alternate days after mRNA delivery; later days are paler. **e**, H929 viability across E:T ratios, as in d. **f**, CAR-positive cells 24 h after electroporation with a cocktail of four CAR mRNAs (BCMA, CD19, HER2 and MUC16 binders), each CAR detected with its own antigen. **g**, CAR density in the same cells. **h**, Surface density of lentivirally transduced FMC63 CD19 CARs; circles, two donors; UTD, untransduced T cells. **i**, IFN-γ during Raji 3D-aggregate co-culture. **j**, Depletion of CD19-positive B cells in PBMCs across E:T ratios. **k**, CAR-positive T cells after CD3-targeted lipid nanoparticles (tLNPs) carrying FMC63 CAR mRNA were added to PBMCs from five donors; mean ± s.d. **l**, B-cell depletion in the same cultures.

To follow expression over time, we transfected primary T cells with `krgr–ζ1–40` or `BBζ` CAR mRNA and measured surface CAR daily for 17 days (Fig. 4b,c). On day 1, `krgr–ζ1–40` T cells displayed 12-fold more CAR than `BBζ` T cells, and, as expected after transient delivery, expression of both CARs then declined at a similar rate. Because it started higher, `krgr–ζ1–40` remained above 2,000 CAR/cell until day 13, compared with day 4 for `BBζ`, and above the 1,000 CAR/cell LLOD through day 17 (Fig. 4b). Thus, after transient delivery, the higher starting expression of `krgr–ζ1–40` kept surface CAR detectable for longer than with `BBζ`, without slowing its decline.

We next asked whether the longer expression of `krgr–ζ1–40` extends its function. Every second day, we co-cultured the CAR T cells with fresh H929 cells and measured IFN-γ and target viability (Fig. 4d,e). `krgr–ζ1–40` T cells released 3.1-fold more IFN-γ than `BBζ` T cells on day 1, and IFN-γ declined over time for both CARs (Figs. 4d and S4a). However, `krgr–ζ1–40` T cells still released IFN-γ on day 13, whereas IFN-γ from `BBζ` T cells fell below the detection limit from day 7. Similarly, at the highest E:T, `krgr–ζ1–40` T cells killed more than 90% of H929 cells through day 13, whereas `BBζ` T cells had lost most of their killing by day 9 (Fig. 4e). From day 1 to day 7, the killing IC50 rose 11-fold for `BBζ` T cells but only 2.5-fold for `krgr–ζ1–40` T cells, which still killed half of the H929 cells through day 13 (Fig. S4b). Together, these results show that `krgr–ζ1–40` extends both CAR expression and function after transient mRNA delivery.

Because `krgr–ζ1–40` yielded more receptor per unit of mRNA (Fig. 4a), we next asked whether it supports the co-expression of several CARs from a shared mRNA dose (Fig. 4f,g). We electroporated T cells with a cocktail of four CAR mRNAs carrying BCMA, CD19, HER2 or MUC16 binders on either `krgr–ζ1–40` or `BBζ`, each at half the single-CAR dose, and detected each CAR with its own antigen in a separate well. With `krgr–ζ1–40`, more than 80% of cells were positive for each CAR, compared with at most a third with `BBζ` (Fig. 4f), and CAR density was 2.6- to 12.9-fold higher (Fig. 4g). Each CAR delivered alone at the full dose showed a similar advantage, with 4.5- to 10.1-fold higher density (Fig. S5). These results show that `krgr–ζ1–40` supports the co-expression of four CARs from a single electroporation at half the per-CAR mRNA dose.

To determine whether these advantages extend to persistent function, we transduced T cells with lentiviral vectors encoding FMC63 anti-CD19 CARs (Fig. 4h,i). `krgr–ζ1–40` T cells displayed 6.8-fold more CAR than `BBζ` T cells (n = 2 donors; Fig. 4h). We then co-cultured the T cells with Raji lymphoma multilayer 3D aggregates at low E:T ratios and measured IFN-γ on days 1, 3, 5 and 7 (Fig. 4i). IFN-γ from both CARs peaked on day 3 and then declined steadily through day 7. `krgr–ζ1–40` T cells released more IFN-γ than `BBζ` T cells in 30 of 32 matched E:T and day conditions. By day 7, when both curves had reached the same plateau, `krgr–ζ1–40` T cells reached half-maximal IFN-γ at a 2.9-fold lower E:T than `BBζ` T cells. Because each T cell must engage more tumor cells in succession at a lower E:T, this shift suggests that `krgr–ζ1–40` T cells stayed active through 2.9-fold more consecutive encounters with antigen-positive tumor cells.

We next asked whether lentivirally transduced `krgr–ζ1–40` T cells also kill target cells more efficiently. In a separate experiment, `krgr–ζ1–40` T cells killed primary human B cells at 2.4-fold lower IC50 E:T than `BBζ` T cells (Fig. 4j), and NALM6 leukemia cells at 1.3-fold lower IC50 E:T (Fig. S4c). Thus, `krgr–ζ1–40` T cells killed both normal and leukemic CD19-positive cells at lower E:T.

Finally, we asked whether `krgr–ζ1–40` enhances CAR T-cell function when the CAR is delivered directly to unselected blood cells [66]. We added CD3-targeted lipid nanoparticles (tLNPs) carrying CAR mRNA to whole PBMC cultures from five donors, so that CAR T cells were generated in situ alongside autologous B cells (Fig. 4k,l). One day later, `krgr–ζ1–40` gave more CAR-positive T cells than `BBζ` at every dose from 12 ng mRNA per 10^6^ cells upward (Fig. 4k). B-cell depletion increased with dose for both CARs, but `krgr–ζ1–40` reached 50% depletion at a 4.6-fold lower dose than `BBζ` (Fig. 4l). Together, these results show that `krgr–ζ1–40` increases CAR expression and enhances CAR T-cell function after mRNA, lentiviral and lipid-nanoparticle delivery.

### Iterative experimental feedback guides CAR ICD refactoring

Having established `krgr–ζ1–40` as the lead compact ICD, we asked whether it could be refined further. In round 3, we worked with AI agents to design 164 variants of `krgr–ζ1–40` and screened them in primary T cells (Fig. S6a,b). The variants tested three groups of changes: 113 rational edits, 30 single and combined point mutations, and 21 synonymous codon recodes. Rational edits changed the juxtamembrane (JM) segment, the costimulatory motif, spacers, charge and length; point mutations tested local sequence changes (Fig. 5a); and codon recodes tested whether expression could be raised through the mRNA sequence alone (Fig. S6e). Surface CAR varied widely among the 21 recodes, which encode the same protein, so codon choice alone changed CAR expression. Of the 122 ICD variants that passed the expression threshold, 17 released at least as much IFN-γ as `krgr–ζ1–40` on the same plate (Fig. S6b).

**Figure 5.**
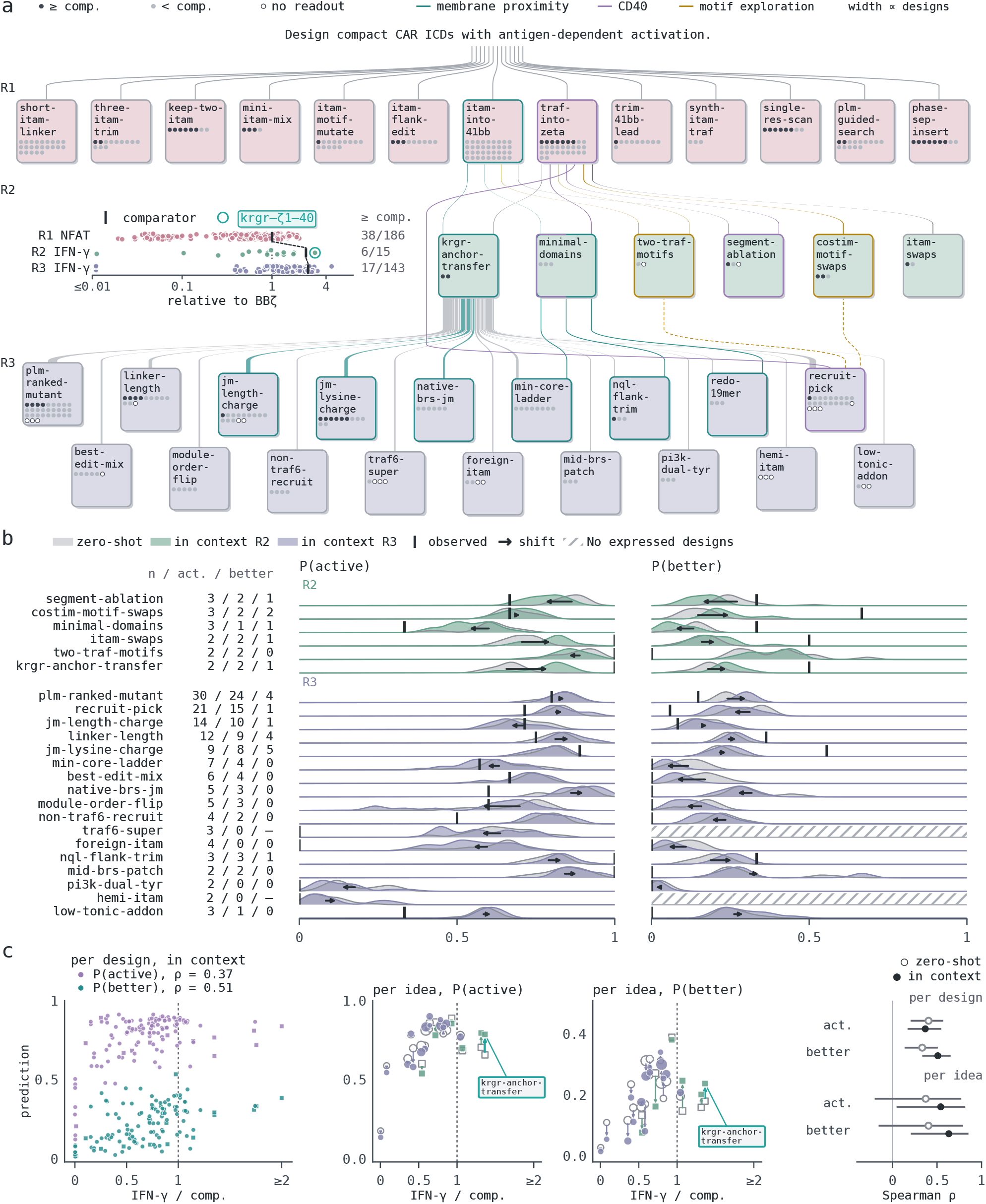
Iterative experimental feedback guides the design of compact ICDs. **a**, Ideas across the three design rounds (left) and each round’s readouts (right). Each card is one idea, labeled by its code name (ideas and code names were generated by AI agents; Supplementary Table 1); dots, its designs (dark, at or above the comparator: `BBζ` (`4-1BB–CD3ζ`) in round 1, the same-plate parent in round 2 and round 3). Ribbons trace lineage; colored borders mark two threads: teal, the distance of the first ITAM tyrosine (Y1) from the membrane; lavender, the CD40 core. Right, every design relative to `BBζ` on its plate, colored by round; black bars, key comparators. **b**, For each idea, the mean predicted probability over its designs that a design is active, P(active), or exceeds the round’s comparator, P(better), from five LLMs (three runs each), zero-shot (gray) or given the earlier rounds’ results (round 2, green; round 3, purple), offsets removed; ridges, distribution over models and runs; arrow, shift of the mean from zero-shot to in context; tick, observed fraction. Left, designs, active designs and designs above the comparator. **c**, Both predictions against IFN-γ relative to the round’s comparator (round 2, `BBζ`; round 3, `krgr–ζ1–40`; dashed line, equal to the comparator) for the 125 designs scored in both tasks; ensemble of the five models with equal weight, each averaged over three runs. Left, per design, in context; squares, round 2; circles, round 3. Middle, per idea, arrow from the zero-shot mean (open) to the in-context mean (filled); size, designs per idea. Right, Spearman correlation with the readout, zero-shot (open) and in context (filled), with bootstrap 95% intervals.

Compact ICDs are difficult to design from structure or from training data. Signaling ICDs such as the CD3ζ cytoplasmic domain are intrinsically disordered [36], so apart from short motifs bound to their partners, such as a phosphorylated ITAM on the ZAP-70 tandem SH2 domains (PDB 2OQ1) [95] and a CD40 peptide on TRAF6 (PDB 1LB6) [89], they offer little structure to model. Their function is measured mainly in cell-based assays, and published CAR libraries offer little training data for compact ICDs [17–24]. Machine learning offers several routes to interpretable sequence design, including libraries of CAR signaling motifs [17], sparse features of protein language models that map to biological concepts [96,97], and grammars that constrain sequence generation [98]. However, these approaches rely on training data or established rules that remain scarce for ICDs. Design therefore relies on reasoning about experimental data, which can be biased toward the designer’s prior beliefs when many choices are defensible [99–103].

To design in this space, we worked with many AI agents in parallel [38,39], which searched the literature and analyzed our assay data. In each round, we and the agents reviewed the preceding measurements together, proposed ideas and developed designs with matched controls. Within each idea, designs changed one feature at a time against matched controls, such as JM charge at fixed length or stepwise trims of ICD length (Fig. S7). The agents wrote and ran the analysis code, and each proposal came with a written rationale [103]. Each design was assigned to an idea when it was proposed, and we traced these ideas across rounds 1–3 (Fig. 5a). Round 1 comprised 186 designs in 14 ideas, and `krgr–ζ1–40` became the parent of the 164 round 3 variants.

Current models did not reliably identify which compact ICDs would fail (Fig. S8). Neither Boltz-2 [104] co-fold confidence nor protein language-model likelihoods (ESM-2 [105] and ProtGPT2 [106]) reliably separated active from inactive ICDs. We also asked five general-purpose LLMs, three queried in a single call and two run as coding agents, given the ICD sequences and a description of the assays, for two probabilities per design: that it was active, P(active), and that it would exceed the round’s comparator, P(better). Each model answered zero-shot and again with the earlier rounds’ results in the prompt (in context). The LLMs called most designs active, including most inactive round 3 variants: the specificity of their calls was 0.15–0.34 zero-shot and 0.18–0.51 in context (Fig. S8a), consistent with the poor calibration of LLM predictions of experimental outcomes [107].

As a retrospective proxy for how much signal the earlier rounds’ data carried about each idea, we measured how far the models’ predictions moved when those data were added to the prompt, a belief change analogous to Bayesian surprise [108,109] (Fig. 5b). Each model was run three times per condition. We removed each model’s overall tendency to call designs active or better, so that ideas were compared by relative belief. Given round 1, the models moved `krgr–ζ1–40` from sixth to third among the 15 round 2 designs for P(active) and from fifth to second for P(better); in the primary T-cell screen, which tested the 15 round 2 designs alongside 78 round 1 designs, it released the most IFN-γ of the 93 ICDs (Fig. 3b). Across ideas, the in-context shift followed the outcome. It correlated with the gap between the observed fraction and the zero-shot prediction (Spearman ρ, 0.52 for P(active) and 0.35 for P(better)), and of the ideas whose prediction moved by 0.03 or more, 12 of 16 moved toward the outcome for P(active) and 11 of 14 for P(better). For 3 ideas in each task, however, the outcome lay outside the in-context range: the models overrated `native-brs-jm`, `foreign-itam` and `traf6-super` for P(active), overrated `plm-ranked-mutant` and `recruit-pick` for P(better) and underrated `jm-lysine-charge`.

To ask what the LLMs learned in context, we compared both predictions with one readout, IFN-γ relative to each round’s comparator, for the 125 round 2 and 3 designs scored in both tasks (Fig. 5c). The earlier rounds’ data raised the correlation of P(better) with the readout for single designs (Spearman ρ, from 0.33 to 0.51) and for the 21 ideas (from 0.40 to 0.63). For P(active), they raised it for ideas (from 0.37 to 0.54) but not for single designs (from 0.40 to 0.37), and in context P(better) tracked single designs more closely than P(active) did (difference in ρ, +0.14; 95% interval +0.01 to +0.28). The size of each design’s update also followed the outcome for P(better) (Spearman ρ, 0.46 across 125 designs; 0.66 in round 2 and 0.44 in round 3): the models raised P(better) for 17 of the 21 designs that exceeded the comparator and for 43 of the 104 that did not. For P(active), the update hardly followed the outcome (ρ, 0.16; Fig. S9b). In both tasks, most of this signal lay between ideas; within ideas, ρ was

0.24 for P(better) and 0.06 for P(active). Thus, the earlier rounds’ data improved the models’ predictions for ideas in both tasks, and for single designs only for P(better).

`krgr–ζ1–40` gave the clearest idea-level update that matched the outcome. It combined two round 1 design ideas: the KRGR anchor of `krgr–ζ1–4-1bb` and the ITAM1 and CD40 core of `brs–ζ1–40` (Fig. 3a). Given round 1, the models raised the predicted activity of this idea more than that of any other round 2 idea (Fig. 5c). In round 3, a single JM edit, R2G (KRGR to KGGR), gave 1.6-fold more surface CAR and 1.2-fold more IFN-γ than `krgr–ζ1–40` in the screen (Fig. S6b) and a 1.5-fold higher IFN-γ peak across E:T ratios (Fig. S6d). This result suggests that further changes to the membrane-proximal JM segment could improve compact ICDs.

We next examined designs for which the earlier rounds gave a clear signal. Round 1 showed that ICDs with Y1 at 8 or less were inactive and that moving Y1 from 5 to 9 with KRGR restored signaling (Fig. S9a). Given the assay descriptions, the models already rated a round 2 design with Y1 at 5 unlikely to be active (mean P(active), 0.16 zero-shot), lowered it further given round 1 (0.10), and it was inactive (Fig. S9b). Round 3 moved this boundary closer to the membrane: a basic JM spacer kept a design active at Y1 = 6, where a neutral spacer did not (Fig. S9a). Basic JM residues contribute to membrane association and signaling [51,110]; in our compact ICDs, JM sequence also influenced how close to the membrane an ITAM could remain functional. Natural ITAM receptors place Y1 at 13–33 residues [111], so with a basic JM segment, compact ICDs can place Y1 closer to the membrane than natural receptors do. Among designs of the same idea, the models’ in-context predictions tracked activity when the designs varied JM length and charge (Spearman ρ, 0.75) and in part when they varied linker length (0.38), but not among the 30 point mutants or the 21 motif swaps (0.01 and −0.19).

The models could also converge on wrong hypotheses. They rated all five designs that restored the native basic-rich JM segment of CD3ζ at about 0.9, yet two were inactive. Although native CD3ζ BRS motifs support membrane association and TCR signaling [52,54], `krgr–ζ1–40` retained antigen-dependent function after most of these segments were removed (Fig. 3b,d,e). Every in-context run called the same 21 of the 43 inactive round 3 designs active, consistent with the correlated errors that LLMs share [112]. Designs that replaced the second YxxL motif of ITAM1 with the CD28 YMNM motif, proposed in several rounds, expressed but did not signal, as ZAP-70 binding requires both YxxL motifs [95,113]. Conversely, the CD40 core, which binds TRAF6 in native CD40 through a motif distinct from those of 4-1BB [88,89], raised IFN-γ in each round: the lead design with the CD40 core released more IFN-γ than the same ICD with the 4-1BB segment in its place (up to 2.4-fold; Fig. S9c), a gain the models did not anticipate (the CD40 design was never predicted more than 0.07 above its counterpart). Asked whether round 3 designs would exceed `krgr–ζ1–40`, the models also overrated the point mutants and motif swaps, of which 4 of 27 and 1 of 17 did, and underrated the JM lysine-charge designs, of which 5 of 9 did (Fig. 5b). These results show that the models followed directions that the earlier data made clear but still missed the gain from the CD40 core and overestimated the promise of several design ideas.

## Discussion

Here we sought to determine how compact a functional CAR ICD can be. We found that most of a clinical second-generation ICD could be removed: `krgr–ζ1–40` retains a single ITAM and a CD40-derived segment in 45 aa. Our results establish three properties of compact ICDs. First, they increase surface CAR expression, across designs, binders and delivery methods. Second, they retain antigen-dependent signaling through defined segments, NFAT through the single ITAM and NF-κB through the CD40-derived motif, with antigen-independent IFN-γ below detection despite higher receptor density. Third, higher expression translated into more durable and broader function after mRNA, lentiviral and lipid-nanoparticle delivery.

Why do compact ICDs express so highly? Because the effect was shared across designs, it may reflect a property of short ICDs more than of a particular sequence. One possibility is that shorter receptors are synthesized, folded or delivered to the cell surface more efficiently. Alternatively, they may be removed from the surface more slowly, because constitutive CD3ζ phosphorylation depends on ICD size [86] and a shorter ICD carries fewer lysines, whose ubiquitination drives CAR downmodulation after antigen engagement [114]. The effect also depended on the binder, as `brs–ζ1–40` did not increase the expression of the FMC63 CAR whereas `krgr–ζ1–4-1bb` did (Fig. 2d). Measuring CAR mRNA, protein synthesis and surface half-life across ICDs and binders would distinguish faster production from slower removal.

Do compact ICDs signal more per receptor, or do they place more receptors on the cell surface? Higher expression contributes: `brs–ζ1–40` lost its IFN-γ advantage with FMC63, where it also lost its expression advantage (Fig. 2d,e), and the NFAT and NF-κB signals of `krgr–ζ1–40` rose in proportion to its surface CAR (Figs. 3e and S2). Signaling per receptor also bears on antigen sensitivity. Because additional ITAMs raise the efficiency with which ligand binding is converted into a signal [15], and CARs are already far less sensitive to antigen than TCRs [16,115], a single-ITAM ICD might need more engaged receptors for the same response. However, when we titrated antigen density [60] at a matched mRNA dose, `krgr–ζ1–40` T cells needed about 1.7-fold more BCMA per target cell than `BBζ` T cells for half-maximal antigen-specific IFN-γ, and released more antigen-specific IFN-γ from a lower antigen-independent floor (Fig. S10). Because `krgr–ζ1–40` displayed 8.4-fold more CAR at this dose (Fig. 4a), its higher surface density likely offsets a lower antigen sensitivity per receptor. Measuring ITAM and ZAP-70 phosphorylation in T cells matched for surface CAR would test this directly.

Our results also bear on the two models that implicate the BRS of the disordered CD3ζ tail in receptor signaling. `krgr–ζ1–40` removed most BRS regions from `BBζ` yet retained antigen-dependent activation (Fig. 3b,d,e). Its antigen-independent IFN-γ also remained below detection at 6.2-fold the surface CAR of `BBζ` (Fig. 3b). Losing most of the BRS therefore impaired neither activation nor antigen dependence, and neither model’s prediction held. For condensation, LAT crosslinking [40,42,116] does not address the result, because LAT condenses after and apart from the TCR–ZAP-70 domain [44,45], and CARs can form microclusters without LAT [117]. Instead, the result runs counter to extending the stickers-and-spacers model of IDR phase separation [46–48] to the ICD, in which losing charged stickers such as the BRS should impair activation. One possibility is that antigen binding and the multivalent partners of the ITAM and CD40 motif, ZAP-70 [95] and TRAF6 [89], cluster the receptor without its disordered tail. For membrane sequestration, in which the BRS hold ITAM tyrosines at the membrane until antigen engagement [50–56], the KRGR anchor and the basic residues after the ITAM may suffice to hold a single ITAM there (Fig. S11). Alternatively, antigen-independent CAR signaling may arise mainly from the 4-1BB domain [118] and additional ITAMs that compact ICDs lack, or from charged patches on the binder [119].

Although the native BRS was dispensable, a short basic segment near the membrane was required. With a basic JM segment, a single ITAM remained functional closer to the membrane than in natural ITAM receptors or `BBζ` (Figs. S9a and S12), and JM segments without lysine or arginine lowered surface CAR. Natural spacing may instead serve regulation, such as holding ITAMs at the membrane until antigen engagement [120]. KRGR, the first four residues of 4-1BB, supplies such a segment. A conserved basic juxtamembrane motif anchors the OX40 tail at the membrane by binding anionic lipids, with similar evidence for 4-1BB and CD40 [110] and for CD28 [121], but to our knowledge KRGR had not been used before as a standalone anchor. Because we measured neither condensates nor basal ITAM phosphorylation, these results argue against a requirement for the BRS rather than against condensation or membrane sequestration. Imaging CAR and LAT clusters separately and measuring ITAM phosphorylation across ICDs that differ in BRS content and juxtamembrane spacing, at matched surface density, would distinguish these possibilities.

How did biological reasoning guide the refactoring? The round 2 designs illustrate the process: measurements from earlier hits motivated combining a short membrane-proximal anchor with an ITAM and a CD40-derived segment. Agents helped turn these observations into sequence proposals, and subsequent experiments determined which combinations worked. Round 3 then tested which changes to the lead preserved or impaired function, allowing us to trace ideas across designs and rounds (Fig. 5a). Because current models did not reliably predict which designs would work (Figs. S8 and S11a–e), we relied on the screens to choose among designs. Experimental feedback guides design in a similar way in directed evolution, antibody design and agent-guided experimentation [122–126]. Given the earlier rounds’ results, the LLMs’ predictions tracked which ideas worked, and tracked which designs worked only when the models were asked whether a design would exceed the comparator (Fig. 5c), consistent with LLMs judging candidates more reliably against one another than on an absolute scale [127]. Whether agent-guided design yields better designs per round than expert or model-based design, and whether it reduces the number of rounds needed, remain to be tested against a baseline matched for experimental budget and data [128], which could also separate the contributions of literature retrieval, biological reasoning and earlier measurements.

Several aspects of our work merit further investigation. The Y1 and JM constraints identified here require prospective tests on new designs. Several experiments used T cells from few donors. Our successive-day challenges measured sustained function rather than serial killing by the same cells, and we did not measure exhaustion, which tonic signaling promotes [11] and rest can reverse [129]. Whether the CD40-derived segment recruits TRAF6 in a CAR, how many cells co-express all four CARs and how compact-ICD CAR T cells perform in vivo remain to be determined.

In summary, successive rounds of agent-guided rational design and experimental feedback identified a 45-aa CAR ICD that increases expression without detectable antigen-independent IFN-γ, extends function after transient mRNA delivery and enhances function after lentiviral and lipid-nanoparticle delivery. The screens also exposed sequence constraints of compact ICDs, such as how close to the membrane the first ITAM tyrosine can sit. Because `krgr–ζ1–40` yielded more receptor per unit of mRNA and occupies less of the payload, compact ICDs may lower the dose needed to generate CAR T cells in vivo [64–67] and enable more complex CAR therapies, such as CAR cocktails.

## Methods

### CAR constructs and mRNA synthesis

Each CAR comprised the CD8α signal peptide, a FLAG (DYKDDDDK) tag, the antigen-binding domain, the CD8α hinge and transmembrane domain, and the ICD. The BCMA binders were the D-domain binder ddBCMA [87] in the Jurkat screen and Fig. 2i, the bb2121 scFv [92] in primary T cells, and the biparatopic VHH of ciltacabtagene autoleucel in Fig. 2i. CD19, HER2 and MUC16 CARs carried the FMC63 scFv [93], a trastuzumab-derived scFv and an anti-MUC16 scFv from Generate Biomedicines, respectively. ICD coding sequences were codon-optimized for human expression with CodonTransformer [130] and DNA Chisel [131]. DNA templates were synthesized as linear double-stranded DNA by Integrated DNA Technologies, as eBlocks for the compact-ICD CARs with the ddBCMA binder and as gBlocks for all other constructs, and were transcribed without linearization. The constructs carried proprietary in-house 5^′^ and 3^′^ UTRs.

mRNA was transcribed in vitro with the HiScribe T7 ARCA mRNA Kit (with tailing) (New England Biolabs, E2060S), which yields a Cap0 structure, and polyadenylated for 1 h with *E. coli* poly(A) polymerase (New England Biolabs, M0276L). Reactions were incubated in a Mastercycler X50 thermocycler (Eppendorf, 6313000018). mRNA was purified with the Total RNA Purification 96-Well Kit (Norgen Biotek, 25100), quantified on a Lunatic spectrophotometer (Unchained Labs) and stored at −80 °C until use.

### Cell lines

Jurkat cells carrying firefly luciferase under an NFAT- or NF-κB-responsive promoter (BPS Bioscience, 60621 and 60651, respectively) were cultured in RPMI 1640 Medium, GlutaMAX Supplement, HEPES (Gibco, 72400047) with 10% fetal bovine serum (FBS; Gibco, A5670502) and 1% penicillin–streptomycin (Gibco, 15140122). HEK293T cells (ATCC, CRL-3216) were cultured in DMEM, high glucose (Gibco, 11965118) with 10% FBS and 1% penicillin– streptomycin. NCI-H929 multiple myeloma cells expressing GFP and firefly luciferase (ATCC, CRL-3580-GFP-LUC2) were cultured in RPMI-1640 with 10% FBS, 0.05 mM 2-mercaptoethanol and 1% penicillin–streptomycin. NALM6 leukemia cells (ATCC, CRL-3273), which we transduced with firefly luciferase lentivirus (BPS Bioscience, 79692-P), and Raji lymphoma cells (ATCC, CCL-86) were cultured in RPMI-1640 with 10% FBS and 1% penicillin– streptomycin. All cells were maintained at 37 °C with 5% CO_2_ and tested negative for mycoplasma by h-IMPACT 1 PCR testing (IDEXX BioAnalytics, 41-00093).

### Primary T cells

Cryopreserved peripheral blood mononuclear cells (PBMCs) from healthy donors were purchased from STEMCELL Technologies (70025). PBMCs were thawed into LymphoONE T-Cell Expansion Xeno-Free Medium (Takara Bio, WK552S) supplemented with 10% human AB serum (MP Biomedicals, 092938249), 10 ng/mL IL-7 (STEMCELL Technologies, 78196.1) and 5 ng/mL IL-15 (STEMCELL Technologies, 78218.1), seeded at 1 × 10^6^ cells/mL and activated with T Cell TransAct (Miltenyi Biotec, 130-111-160) at 1:100. Cells were counted with acridine orange/propidium iodide (AO/PI) staining (Revvity, CS2-0106) on a Cellometer K2 (Revvity), and cells with post-thaw viability below 75% were not used. T cells were expanded for three to five days before receptor delivery.

### mRNA electroporation

T cells were washed twice in Opti-MEM Reduced Serum Medium (Gibco, 31985070) and resuspended in Opti-MEM at 1 × 10^6^ cells per 20 µL well of the 96-well Nucleocuvette plates from the P3 Primary Cell 4D-Nucleofector X Kit (96-well) (Lonza, V4SP-3096). mRNA was added at 4 µg per 10^6^ cells unless stated, and cells were electroporated on a 4D-Nucleofector with the 96-well Shuttle (Lonza) using pulse code DS-137. Cells were recovered immediately into prewarmed medium with 10% human serum at 1 × 10^6^ cells/mL and rested overnight. Jurkat reporter cells were electroporated with the SF Cell Line 4D-Nucleofector X Kit (96-well) (Lonza, V4SC-2096) using pulse code CL-120, with 5 × 10^5^ cells and 8 µg of mRNA per well.

### Lentiviral transduction

Lentiviral particles encoding FMC63 CARs with the `BBζ` or `krgr–ζ1–40` ICD were custom-produced by Vector-Builder. PBMCs were plated at 1 × 10^5^ cells per well in Falcon 96-well clear flat-bottom TC-treated plates (Corning, 353072) and activated with TransAct. After 24 h, lentiviral particles were added directly to the cells at a multiplicity of infection of 10, in T-cell medium without human serum and without a transduction enhancer or spinoculation. On day 4, cells were pooled into G-Rex 6-well plates (Wilson Wolf, 80240M) and expanded until 10 days after transduction, when CAR density was measured and the co-culture assays were set up. Untransduced T cells from the same donor served as controls.

### CD3-targeted lipid nanoparticles

CD3-targeted lipid nanoparticles (tLNPs) encapsulating FMC63 CAR mRNA with the `BBζ` or `krgr–ζ1–40` ICD were produced by GenScript (lots U236FPBQG0-1B and U9797RPHG0-5A, respectively), with an anti-CD3 antibody (clone OKT3) as the targeting ligand and SM-102 as the ionizable lipid. The encapsulated mRNA carried GenScript UTRs, a Cap1 structure, N1-methyl-pseudouridine and a 100-nt poly(A) tail. PBMCs were thawed without activation into LymphoONE with 10% human serum and cultured at 25 × 10^6^ cells per 30 mL in G-Rex 6-well plates. One day after thaw, tLNPs were added directly to the PBMC cultures in an eight-point, 3-fold dilution series starting at 1 µg mRNA per 10^6^ cells. Cells were analyzed by flow cytometry one day later. Because tLNPs were added to whole PBMCs, CAR T cells were generated in the same culture as the autologous B cells they targeted.

### Flow cytometry and CAR quantification

Cells were stained in BSA Stain Buffer (BD Biosciences, 554657) with a PE-conjugated anti-DYKDDDDK antibody, REAfinity (Miltenyi Biotec, 130-128-816), at 1:100 for 1 h at 4 °C in Falcon 96-well non-treated V-bottom plates (Corning, 353263). FMC63 CARs, including those in Fig. 2d, were detected with the same antibody. Cells were acquired on an ID7000 Spectral Cell Analyzer (Sony Biotechnology) together with QuantiBRITE PE beads (BD Biosciences, 340495) run in the same session at the same settings. CAR molecules per cell were interpolated from the bead standard curve of PE molecules per bead, assuming one PE molecule per antibody. Lymphocytes were gated on forward against side scatter, and CAR-positive cells on PE fluorescence above a CAR-negative reference (Fig. S3). We report the CAR-positive percentage and CAR molecules per CAR-positive cell. The lower limit of detection was 1,000 CAR molecules per cell. Flow data were analyzed in a gating viewer that we built in-house as part of our electronic lab notebook.

### Jurkat reporter assays

To make BCMA-positive targets, we transfected HEK293T cells at 100,000 cells per well with 2.4 µg of linear DNA encoding full-length BCMA using Lipofectamine 3000 Transfection Reagent (Invitrogen, L3000001) and plated them at 60,000 cells per well in Nunc F96 MicroWell white polystyrene plates (Thermo Scientific, 236105). BCMA expression was confirmed by flow cytometry about 24 h after transfection with a PE-conjugated anti-CD269 (BCMA) antibody, REAfinity (clone REA315; Miltenyi Biotec, 130-118-970), and untransfected HEK293T cells served as BCMA-negative targets. CAR-expressing reporter cells were added 18 h after electroporation at a 1:1 ratio of CAR-positive cells to target cells and co-cultured for 4 h at 37 °C. Luminescence was developed in the same wells with 35 µL of ONE-Glo Luciferase Assay System (Promega, E6120) and read on an EnVision Multimode Plate Reader (Revvity).

The library screen (Fig. 1c,d) comprised two independent experiments. For each design, we divided its mean luminescence across the two experiments by the mean of the `BBζ` wells in the same condition. In the KRGR comparison (Fig. S1b), each BCMA-stimulated well was divided by the same-plate `BBζ` well. For the signaling mutants (Figs. 2h and 3e), NFAT and NF-κB reporter cells were electroporated with the same constructs in parallel and stimulated with BCMA-positive HEK293T cells. We report raw luminescence from two technical readings and, in Fig. S2, luminescence divided by the CAR density measured in the same reporter line.

### Co-culture assays with primary T cells

Luciferase-expressing target cells were plated at 10,000 cells per well, and effector T cells were added at the stated E:T ratios in 100 µL of LymphoONE with 10% human serum and without cytokines, in 96-well solid white flat-bottom TC-treated plates (Corning, 3917). E:T ratios count CAR-positive T cells per target cell unless stated. After 24 h at 37 °C, we collected 50 µL of supernatant for IFN-γ and read residual target-cell luminescence with ONE-Glo in white plates on the EnVision reader. Lower luminescence indicates more killing. Tonic IFN-γ was measured from wells on the same plate that held T cells without target cells.

IFN-γ was measured with the BD CBA Human IFN-γ Flex Set (Bead E7) (BD Biosciences, 558269), with BSA Stain Buffer (BD Biosciences, 554657) as the assay buffer. Supernatant (50 µL) was incubated with 50 µL of capture bead mix and 50 µL of detection reagent for 3 h at room temperature on a plate shaker. Samples were then washed once, acquired on the ID7000 and quantified against a standard curve in our electronic lab notebook. The lower limit of detection was 9.8 pg/mL. Samples above the standard range were diluted.

### Primary T-cell screens

In the primary T-cell re-screen (Figs. 1e and 3b), the 78 round 1 designs, the 15 round 2 designs, `brs–ζ1–40`, `BBζ` and negative controls were expressed as bb2121 CARs in T cells. CAR density was measured 24 h after electroporation, and T cells were co-cultured with H929 cells at 5:1 for 24 h. IFN-γ was measured on two replicate plates (Fig. S1c), and values were divided by those of the same-plate `BBζ`. In the round 3 screen (Fig. S6a,b), each of the 164 `krgr–ζ1–40` variants was electroporated once into T cells and co-cultured with H929 cells at 10:1 in two wells, and IFN-γ was averaged across the two wells. Each of the two plates carried `krgr–ζ1–40`, `BBζ`, ITAM-null and no-mRNA controls, and variant values were divided by those of the same-plate `krgr–ζ1–40`. Variants with fewer than 30% CAR-positive cells were excluded.

### E:T titrations

For the characterization panel (Fig. 2a–g), ten compact ICDs, `BBζ` and a negative control were expressed as bb2121 or FMC63 CARs and co-cultured with H929 or NALM6 cells at eight 2-fold E:T ratios from 20:1 to 0.16:1. BCMA co-cultures had two replicate wells and CD19 co-cultures had one well per condition. For Fig. 3d, round 2 designs including `krgr–ζ1–40`, `brs–ζ1–40` and `BBζ` were co-cultured with H929 cells at twelve 3-fold E:T ratios from 20:1 to 0.0001:1, in two wells per condition. For Fig. S6c,d, `krgr–ζ1–40` and four round 3 hits were co-cultured with H929 cells at twelve 2-fold E:T ratios from 10:1 to 0.0049:1, in two wells per condition. Killing was normalized to each construct’s lowest E:T in Fig. 2c,f, to each construct’s highest well in Fig. 3d and to each construct’s low-E:T plateau in Fig. S6c. For Fig. 2i, `brs–ζ1–40` and `BBζ` CARs with the VHH or ddBCMA binders were measured about 24 h after electroporation, in T cells from a different donor for each binder.

### Expression and function after transient mRNA delivery

For the mRNA dose titration (Fig. 4a), T cells were electroporated with twelve 2-fold doses of CAR mRNA from 4 to 0.002 µg per 10^6^ cells, and CAR density was measured 24 h later. Cells electroporated with the lowest dose of `krgr–ζ1–40` mRNA served as the CAR-negative reference for the 4 µg plots in Fig. S3.

To follow expression and function over time (Figs. 4b–e and S4), we electroporated T cells with 4 µg of `krgr–ζ1–40` or `BBζ` CAR mRNA and cultured them in LymphoONE with 10% human serum, IL-7 and IL-15, refreshing the medium every other day. CAR density and the CAR-positive fraction were measured daily from day 1 to day 17. On every second day from day 1 to day 17, T cells were co-cultured with fresh H929 cells (5,000 per well) for about 24 h, at eleven 2-fold E:T ratios from 20:1 that count total T cells, with one well per condition. H929 viability was normalized to the highest luminescence of each construct and day.

### Multi-CAR co-expression

T cells were electroporated on day 4 after activation with a cocktail of four CAR mRNAs carrying BCMA (bb2121), CD19 (FMC63), HER2 (trastuzumab) and MUC16 binders on the same ICD, 2 µg of each (8 µg in total), or with each CAR mRNA alone at 4 µg (Fig. S5). After 24 h, each CAR was detected in a separate well with its PE-conjugated antigen (ACROBiosystems, BCA-HP2H7 for BCMA, CD9-HP2H5 for CD19 and HE2-HP2E3 for HER2), the MUC16 CAR with biotinylated human MUC16 (residues 13810–14451, Fc, Avitag; ACROBiosystems, CA5-H82F8) and streptavidin–PE (BD Pharmingen, 554061), and all with QuantiBRITE calibration. Untransfected T cells served as the CAR-negative control.

### Antigen-density titration

HEK293T cells (ATCC, CRL-3216) that we transduced with firefly luciferase lentivirus (BPS Bioscience, 79692-P) were transfected at 100,000 cells per well with eight 2-fold amounts of linear DNA encoding full-length BCMA, from 2.4 µg, using Lipofectamine 3000 to give eight BCMA densities, which we measured with the anti-BCMA antibody above and QuantiBRITE PE beads at co-culture setup. T cells electroporated with 4 µg of `krgr–ζ1–40` or `BBζ` CAR mRNA were co-cultured with these targets at 5:1 for 18 h, with one well per condition, and IFN-γ was measured as above (Fig. S10).

### Assays with lentivirally transduced T cells

The CAR density of lentivirally transduced T cells (Fig. 4h) was measured by anti-FLAG staining with QuantiBRITE calibration 10 days after transduction. For the 3D aggregate co-culture (Fig. 4i), Raji cells were seeded at 100,000 cells per well in round-bottom ultra-low-attachment spheroid microplates (Corning, 4520), centrifuged at 200 × g for 10 min and left to aggregate for 72 h. Transduced or untransduced T cells were then added at eight 2-fold E:T ratios from 0.1:1 to 0.00078:1, counting CAR-positive T cells against the seeded Raji cells, with one well per condition. On days 1, 3, 5 and 7, 100 µL of supernatant was collected for IFN-γ and replaced with 100 µL of fresh LymphoONE with 10% human serum and 100 IU/mL IL-2 (Miltenyi Biotec, 130-097-743).

For B-cell depletion (Fig. 4j), autologous PBMCs were seeded to give 10,000 CD3^−^CD19^+^ B cells per well, based on the CD19^+^ fraction measured by flow cytometry before seeding. Transduced T cells, activated and transduced separately from the same PBMCs, were added back at twelve 2-fold ratios of CAR-positive T cells to B cells, from 20:1 to 0.0098:1, in 100 µL of cytokine-free LymphoONE with 10% human serum, with one well per condition. After 24 h at 37 °C, cells were stained with anti-CD3–BV510 (clone UCHT1; BD Biosciences, 563109) and anti-CD19–FITC (clone HIB19; BD Biosciences, 555412), and B cells were counted as CD3^−^CD19^+^ events by flow cytometry, without a viability dye. Each count was expressed as a percentage of the highest count for the same construct, and depletion as 100 minus this percentage. In the same experiment, NALM6-luc cells were plated at 10,000 cells per well and co-cultured with the transduced or mock T cells for 24 h at twelve 2-fold E:T ratios from 20:1. Residual luminescence was read as in the other co-culture assays, and killing was calculated relative to mock T cells at each ratio (Fig. S4c).

### B-cell depletion after tLNP delivery

PBMCs from five donors were dosed with tLNPs as above (Fig. 4k,l). One day later, cells were stained with anti-DYKDDDDK–PE, anti-CD3 and anti-CD19 as above. CAR-positive T cells are reported as a percentage of CD3^+^ cells. B-cell counts were expressed as a percentage of the highest count in the same plate row, and depletion as 100 minus this percentage. Four donors had two replicate rows per ICD, averaged after normalization, and one donor had one.

### Dose–response fitting and replicate agreement

Data were analyzed in Python with NumPy, SciPy and Matplotlib. Killing and B-cell depletion curves were fitted with a decreasing four-parameter logistic, *y* = *B* +*A*/[1 +(*x*/*C*)^*h*^], by unweighted least squares over all normalized readings. The bottom B was bounded between 0 and 100%, the amplitude A between 0 and 200 percentage points, the Hill slope h between 0.1 and 8, and the midpoint C to within two decades of the tested range. We kept the best of 12 starting points. IC50 is the E:T or dose at which the fitted curve crosses 50% viability. IFN-γ against E:T (Fig. 4i) or antigen density (Fig. S10) was fitted with an increasing four-parameter logistic with all four parameters free. Agreement between replicate plates (Fig. S1c,d) is the squared Pearson correlation of log_10_ values for wells with both replicates above the detection limit. Error bars show the mean ± s.d. of replicate wells.

### Agent-guided sequence design

We used AI agents with access to computational tools to propose compact CAR ICDs from scientist-defined objectives and reference sequences. In round 1, we used Claude Opus 4.6, Claude Sonnet 4.6, GPT-5.2, Kimi K2.5 and MiniMax M2.5. `brs–ζ1–40` was proposed by Claude Opus 4.6 in round 1. We prompted the agents to shorten CD3ζ, vary ITAM number and identity, incorporate costimulatory motifs, or jointly redesign activation and costimulatory elements (Supplementary materials, “Round 1 design prompts”).

The agents had access to literature search, protein annotation, sequence analysis, Python execution, structure prediction and visual assessment of predicted structures. The computational tools ran as serverless functions on Modal.

In rounds 2 and 3, we worked with Claude Code (Opus 4.7 and Opus 4.8, respectively) to interpret experimental measurements and refine the designs. We used literature evidence and results from earlier rounds to define design objectives, revise candidate sequences and develop matched controls.

We organized the designs of each round by idea. An idea was a design question with a written rationale, for example whether the KRGR·KQGQ JM segment acts through charge or through spacing. Each design was assigned to one idea when it was proposed, and the agents gave each idea a short code name. Because no screen could cover all the sequences that an idea allows, the designs within an idea were chosen as contrasts that one plate could interpret. Each design changed one feature relative to the same-plate comparator, and the set included charge-matched pairs at fixed length, residue-by-residue trims, re-tests of earlier designs beside matched controls and tyrosine-null calibrators. For some designs, the rationale also stated the expected direction of the result. Fig. S7 shows six ideas organized in this way, including one whose readouts did not separate its designs.

### Evaluation of agent-generated design hypotheses

We evaluated the agents’ design hypotheses retrospectively against the screens (Figs. 5b,c, S8 and S9).

#### Design data

The benchmark comprised 339 unique ICDs with activity calls: 186 from round 1 (Jurkat NFAT; active at 0.25 or more of `BBζ`), 15 from round 2, screened in primary T cells alongside the 78 re-screened round 1 designs, which thus also received round 2 calls (IFN-γ, active at 0.5 or more of the same-plate `BBζ`), and 138 new round 3 ICDs (H929 killing, active at 0.5 or more of `krgr–ζ1–40`). Constructs encoding the same protein took the majority call; plate controls were excluded, as was `krgr–ζ1–40` from round 3, where it served as the reference. For P(better), a design exceeded the comparator when its IFN-γ was at least that of the comparator on the same plate (round 2, `BBζ`, 6 of 15 designs; round 3, `krgr–ζ1–40`, 16 of the 117 designs with IFN-γ).

#### Predictions

Five LLMs, each run three times on the same task text (three queried directly and two run as coding agents), gave the probability that each ICD was active, P(active), and, in a separate task, that its readout exceeded that of the round’s comparator, P(better), zero-shot and in context, that is, given the measured ICDs of the earlier rounds (round 1 for round 2; rounds 1 and 2 for round 3). To compare ideas by relative rather than absolute belief, we removed each model’s offset: for every model, round and condition, one shift on the log-odds scale set the model’s mean prediction over the round’s test ICDs to the active fraction of the earlier rounds’ labeled ICDs (those given in the in-context prompt), so that no test label was used. Because one shift is applied to all of a model’s predictions in a round and condition, it leaves each model’s ranking of the designs, and so its AUROC, unchanged. It removes the models’ overall overprediction of activity (in round 3, a mean zero-shot prediction of 0.78 against an observed active fraction of 0.66), which would otherwise dominate the comparison of zero-shot and in-context predictions. For ensemble predictions, probabilities were averaged over runs within a model, and the five models were combined with equal weight.

#### Metrics

AUROC and specificity were computed per run against the measured calls, with 95% intervals from 1,000 bootstrap resamples of ICDs (Fig. S8). Specificity was computed for each model’s own calls and, independent of its calling threshold, at the highest threshold on its predicted probability that still called at least 90% of active ICDs active, with ties at the threshold called active. For each idea, we compared the mean prediction over its designs, zero-shot and in context, with the fraction of its designs observed active or above the comparator (Fig. 5b). An in-context prediction supported the outcome when its mean lay at least 0.05 closer to the observed fraction than the zero-shot mean. An idea was underrated or overrated when, treating its designs as independent trials with the in-context probabilities, a count as extreme as the one observed had a probability below 0.10, with the observed fraction above or below the in-context mean. The in-context shift of an idea was compared with the gap between its observed fraction and its zero-shot mean by Spearman correlation, and among ideas that moved by 0.03 or more, by the share that moved toward the outcome.

Finally, the Spearman correlation between the ensemble prediction and IFN-γ relative to each round’s comparator was computed per design and per idea (the mean over its designs, with ratios capped at 2), for both predictions on the designs scored in both tasks, with intervals from 2,000 bootstrap resamples of designs or ideas (Fig. 5c). For ideas with at least four designs, the correlation of in-context P(active) with the activity readout was computed among the designs of one idea (Fig. S9b).

## Acknowledgments

We thank Mike Nally, John Ingraham, Molly Gibson, and Kaggen Ausma for their feedback and support.

## Supplementary materials

**Figure S1.**
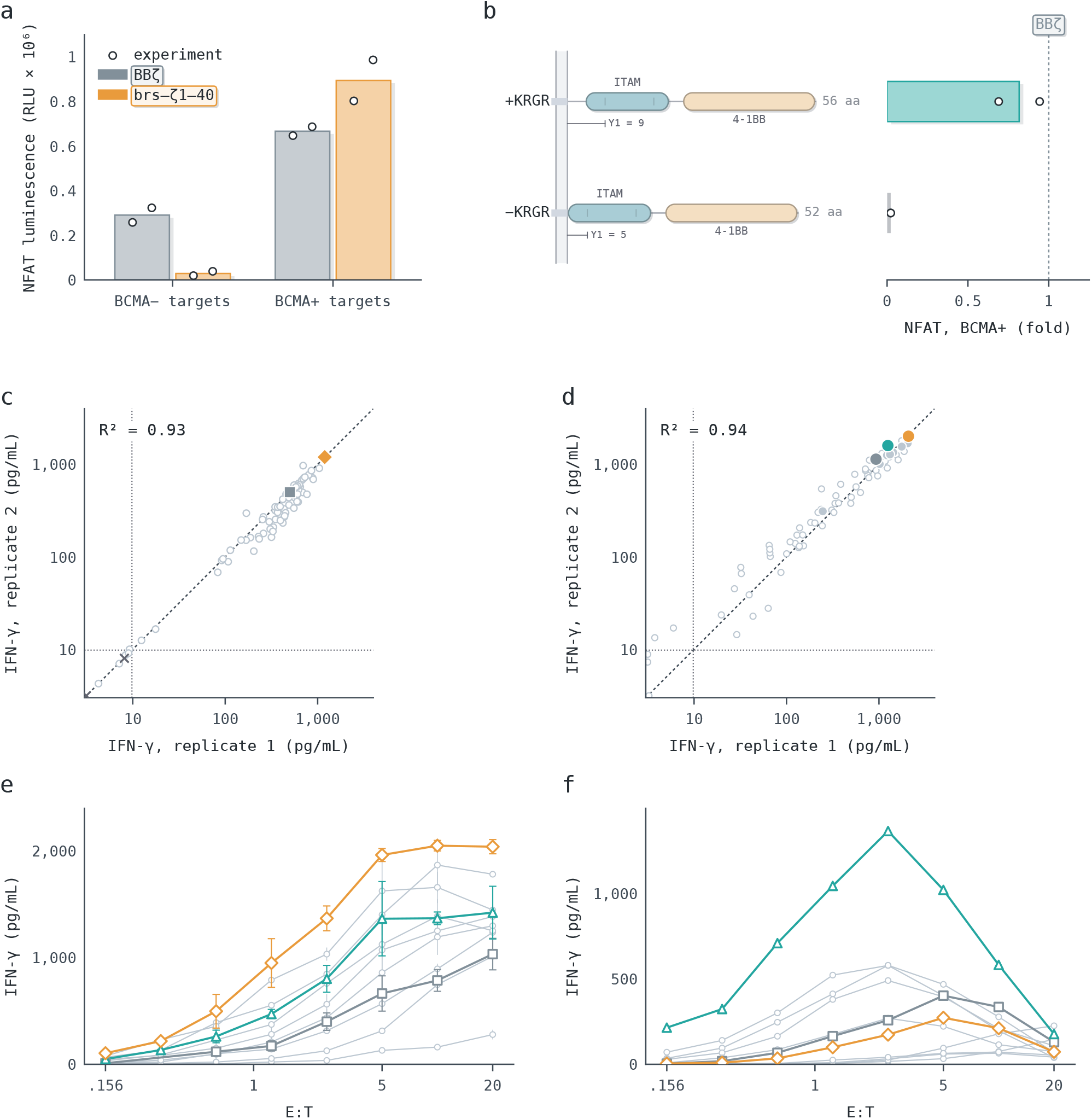
Individual-experiment Jurkat NFAT responses, the KRGR comparison, agreement between replicate IFN-γ measurements and IFN-γ across E:T ratios. **a**, NFAT luminescence of `brs–ζ1–40` and `BBζ` (`4-1BB–CD3ζ`) without antigen and with BCMA-positive targets in each of two independent experiments. **b**, BCMA-stimulated NFAT for `krgr–ζ1–4-1bb` with (+KRGR) and without (−KRGR) its four N-terminal residues, relative to the same-plate `BBζ` (dashed line). Without KRGR, the first ITAM tyrosine (Y1) sits four residues closer to the membrane and the design did not function. **c**,**d**, H929-induced IFN-γ in replicate 1 against replicate 2, measured on separate plates; dashed line, identity; dotted lines, lower limit of detection; R^2^, squared Pearson correlation of log_10_ values for wells above that limit (c, 74 wells; d, 73 wells). **c**, The primary T-cell re-screen of Fig. 1e, which plots replicate 1: library designs (light gray circles), `brs–ζ1–40` (orange diamond), `BBζ` (gray square) and negative controls (×). **d**, The BCMA CARs of Fig. 2b: small open light-gray circles, non-peak E:T wells; filled circles, each design’s peak, as ranked in Fig. 2b: `brs–ζ1–40` (orange), `krgr–ζ1–4-1bb` (teal), `BBζ` (gray) and other designs (small, light gray). **e**,**f**, IFN-γ across the eight E:T ratios behind the peak values of Fig. 2b (e; H929 targets, BCMA CARs; mean ± s.d. of the two replicate plates, or the single reading where only one exists) and Fig. 2e (f; NALM6 targets, CD19 CARs; one well per E:T), with points joined without fitting: `brs–ζ1–40` (orange diamonds), `krgr–ζ1–4-1bb` (teal triangles), `BBζ` (gray squares) and other designs (light gray circles).

**Figure S2.**
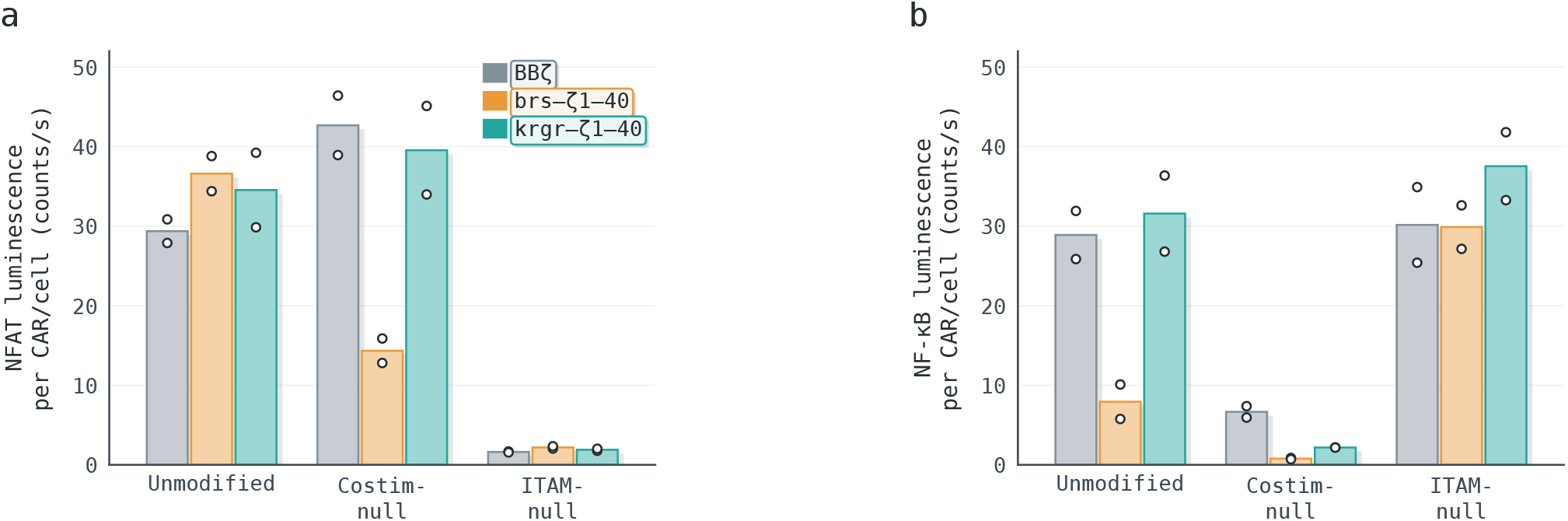
Reporter signal per surface CAR for the signaling mutants of Figs. 2h and 3e. **a**,**b**, Jurkat NFAT (a) and NF-κB (b) luminescence divided by surface CAR density for the unmodified, costimulation-null and ITAM-null forms of `BBζ` (`4-1BB–CD3ζ`), `brs–ζ1–40` and `krgr–ζ1–40`. Bars, means; circles, individual readings.

**Figure S3.**
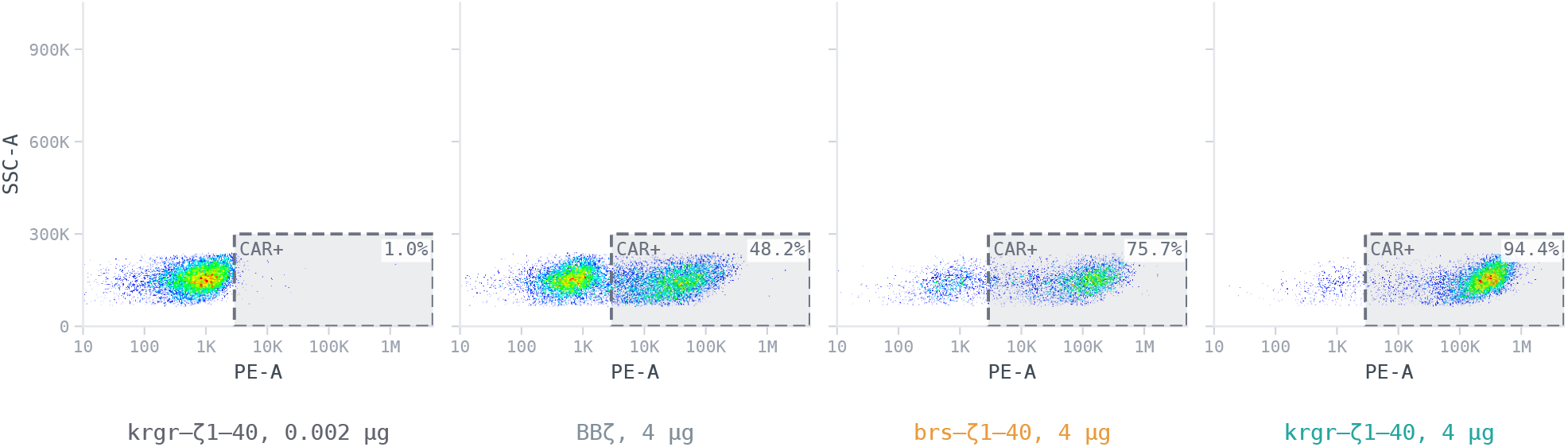
Surface CAR staining at the highest mRNA dose of Fig. 4a. Side scatter against anti-CAR staining of primary T cells expressing `BBζ` (`4-1BB–CD3ζ`), `brs–ζ1–40` or `krgr–ζ1–40` CARs, beside a CAR-negative reference (left). Color shows event density; dashed rectangles, the CAR-positive gate, with the percentage of CAR-positive cells.

**Figure S4.**
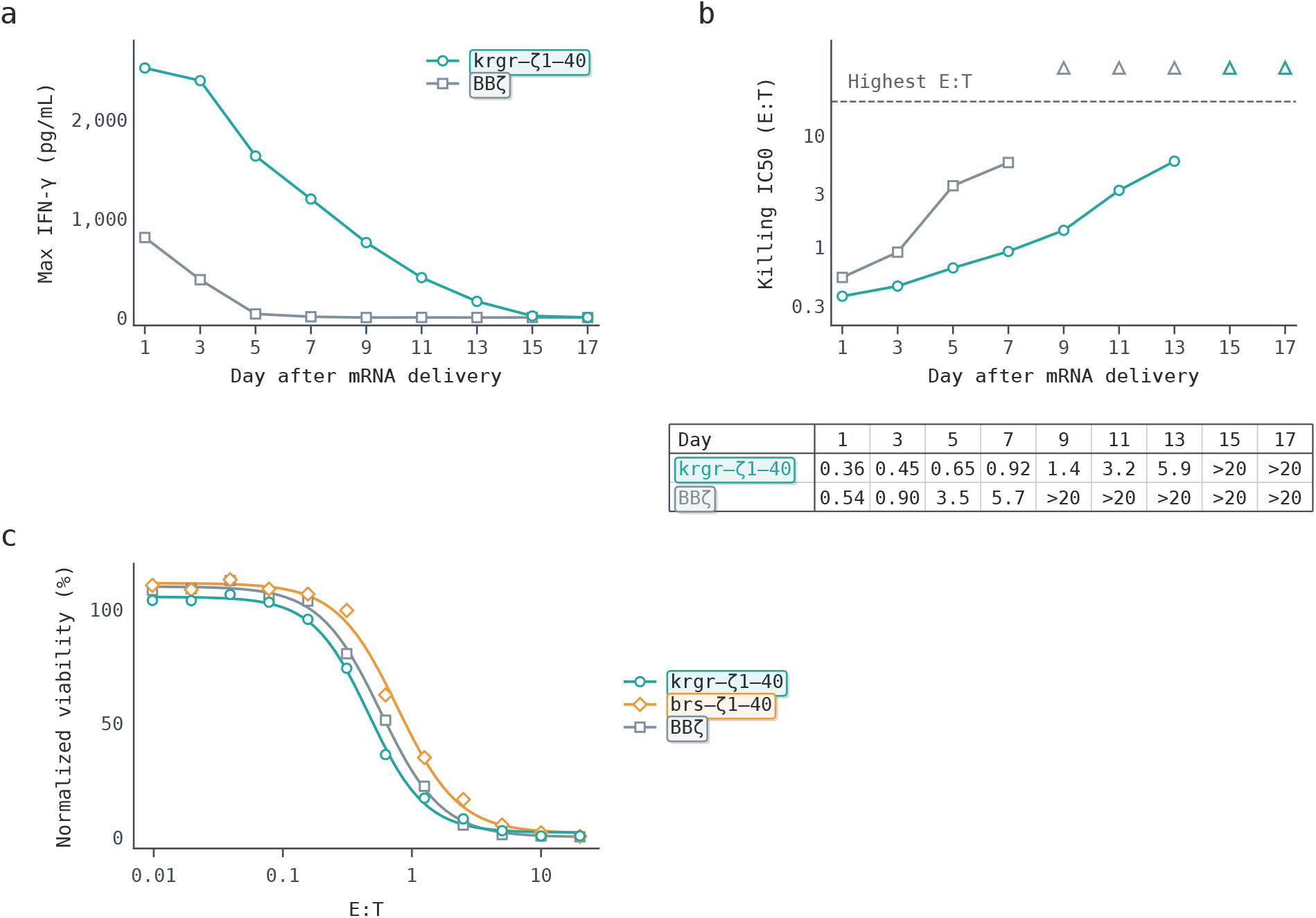
Loss of IFN-γ release and killing after mRNA electroporation, and NALM6 killing by lentiviral CAR T cells. **a**, Highest IFN-γ across the E:T ratios of Fig. 4d on each day. **b**, Killing IC50 on each day, the E:T at which the fitted H929 viability curves of Fig. 4e cross 50%; triangles, days whose fits stay above 50% at the highest E:T. The values under b list the same IC50s (>20, above 50% at the highest E:T). **c**, Viability of NALM6 leukemia cells after co-culture with the lentivirally transduced CAR T cells of Fig. 4j, and with `brs–ζ1–40` CAR T cells from the same experiment, across twelve 2-fold E:T ratios, as 100 minus killing relative to mock T cells; curves, descriptive four-parameter fits as in Fig. 4j.

**Figure S5.**
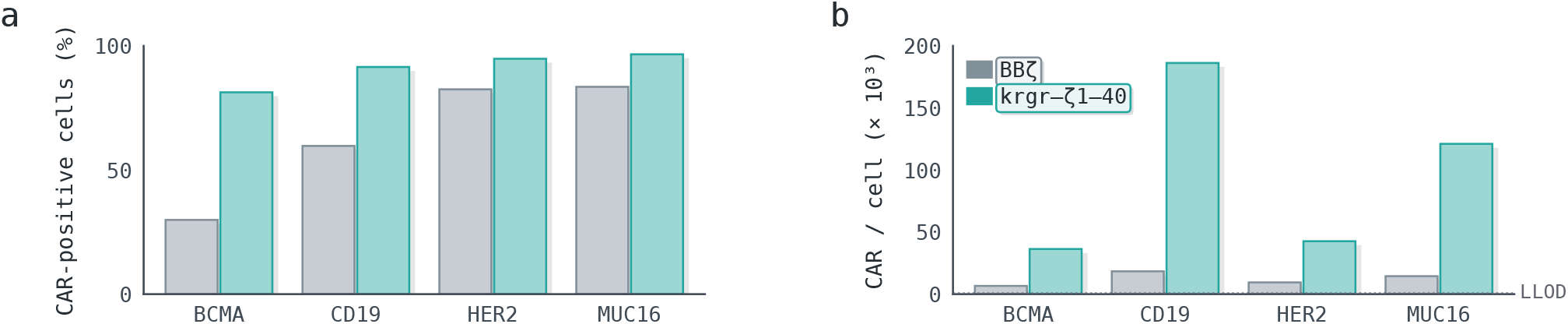
Single-CAR expression after mRNA electroporation. **a**,**b**, Each of the four CARs of Fig. 4f,g (BCMA, CD19, HER2 and MUC16 binders) electroporated alone on `BBζ` (`4-1BB–CD3ζ`) or `krgr–ζ1–40`: CAR-positive cells (a) and CAR molecules per cell (b) at 24 h, each CAR detected with its own antigen. Dotted line, lower limit of detection.

**Figure S6.**
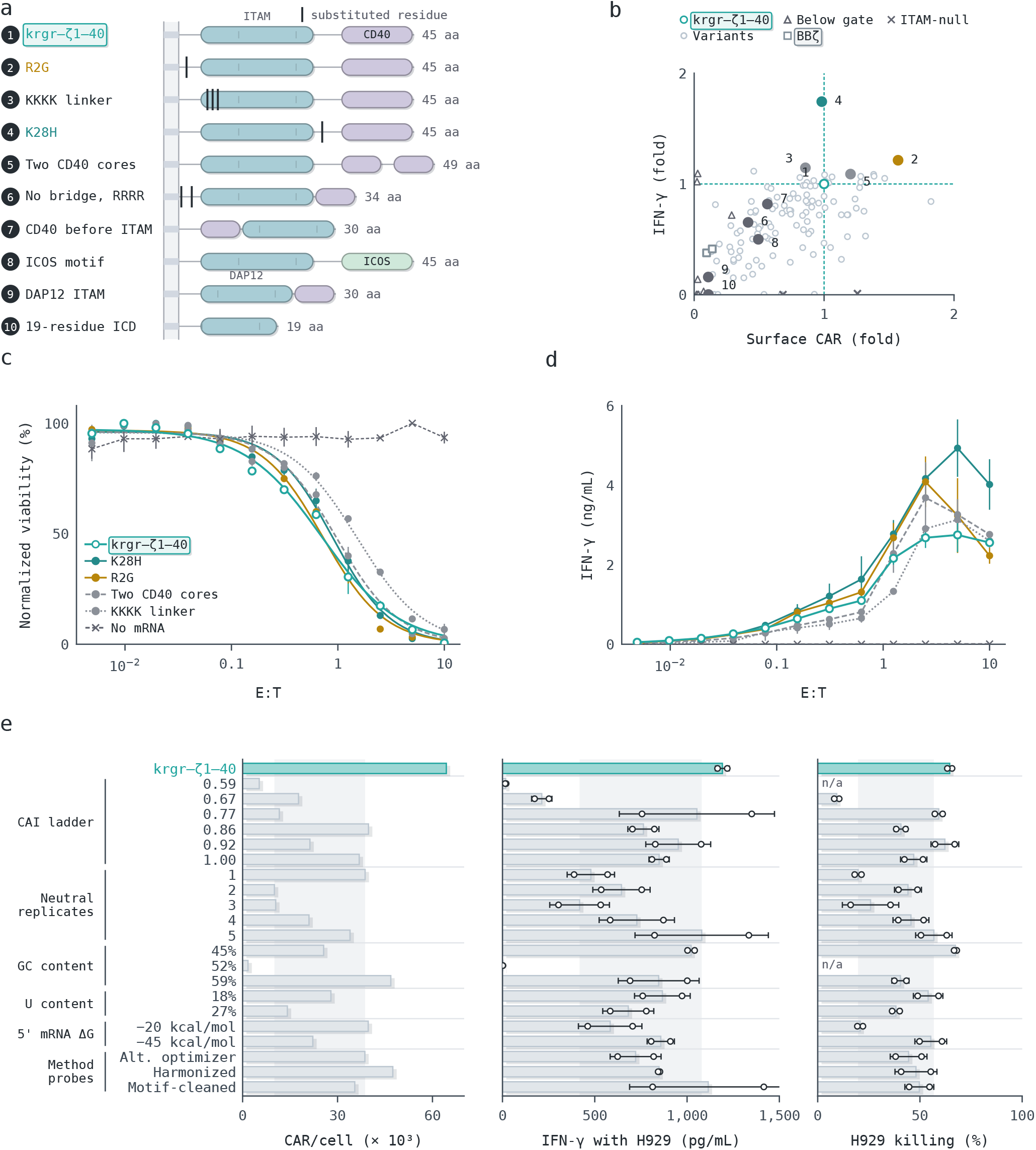
Round 3 variants of krgr–ζ1–40. **a**, Representative round 3 designs drawn to residue scale with segments as in Fig. 1b and numbered as in b; ticks mark substituted residues. **b**, Surface CAR density and H929-induced IFN-γ of each variant, relative to the same-plate `krgr–ζ1–40`; open circles, variants that passed the expression gate; triangles, variants below the gate with a measured CAR density; filled circles, the designs of a. **c**,**d**, H929 viability (c) and IFN-γ (d) across E:T ratios for `krgr–ζ1–40` and four screen hits. **e**, Synonymous codon recodes of the `krgr–ζ1–40` CAR, grouped by codon-design family, with their CAR/cell, H929-induced IFN-γ and killing below `krgr–ζ1–40` with its validated codons; pale band, the range of five neutral recodes; n/a, failed the expression gate.

**Figure S7.**
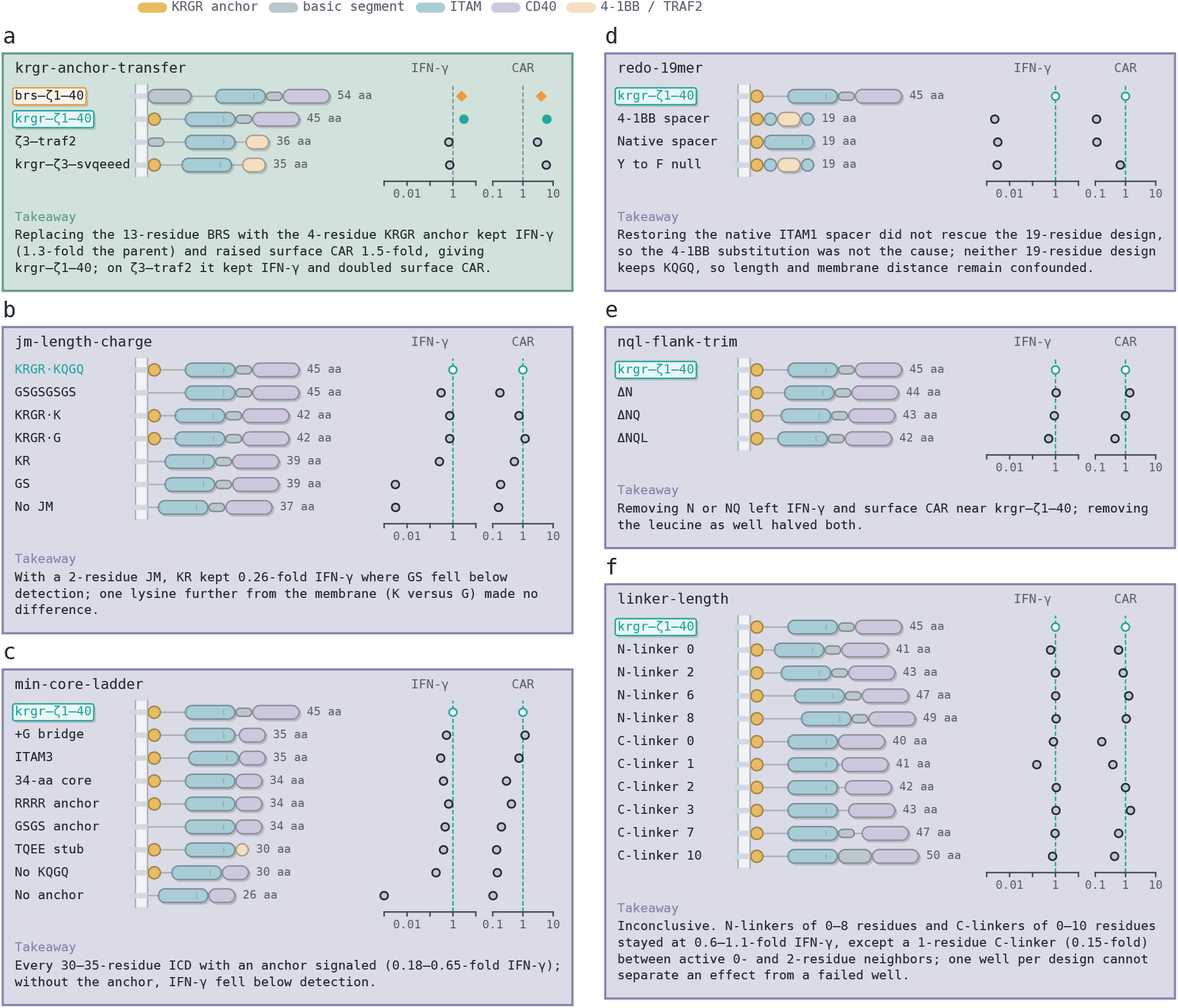
Design ideas built as controlled contrasts. Each card is one idea of Fig. 5a, labeled by its code name and framed in its round’s color (green, round 2; indigo, round 3). Left, the idea’s designs drawn to residue scale from the transmembrane end, with segments as in Fig. 1b; right, IFN-γ and surface CAR relative to the round’s comparator on the same plate (dashed line): `BBζ` (`4-1BB–CD3ζ`) in round 2 and `krgr–ζ1–40` in round 3, which is drawn as an open teal circle. **a**, Transfer of the KRGR anchor onto two round 1 hits, `brs–ζ1–40` (orange diamonds) and `ζ3–traf2`. **b**, JM length and charge, with charge-matched pairs at fixed length. **c**, Shorter ICDs that keep ITAM1 and the CD40 core. **d**, The inactive 19-aa design re-tested beside a matched-length design with the native ITAM1 spacer and a tyrosine-null calibrator. **e**, Stepwise trimming of the three residues before the first ITAM1 tyrosine. **f**, An inconclusive idea: the length of the linkers before (N-linker) and after (C-linker) ITAM1. One well per round 3 design, each relative to `krgr–ζ1–40` on its own plate.

### LLM evaluations of designs

#### Data

The benchmark uses 339 unique ICD proteins with activity calls from three rounds: round 1, the library screen (186 ICDs; Jurkat NFAT, active at 0.25 or more of `BBζ`); round 2, the 15 new designs and 78 re-screened round 1 designs in primary T cells (93 ICDs; IFN-γ, active at 0.5 or more of the `BBζ` CAR on the plate); and round 3, the `krgr–ζ1–40` variants of Figs. 5 and S6 (138 ICDs not measured before; H929 killing, active at 0.5 or more of `krgr–ζ1–40`). Constructs encoding the same protein take the majority call; plate controls and `krgr–ζ1–40` are excluded.

**Figure S8.**
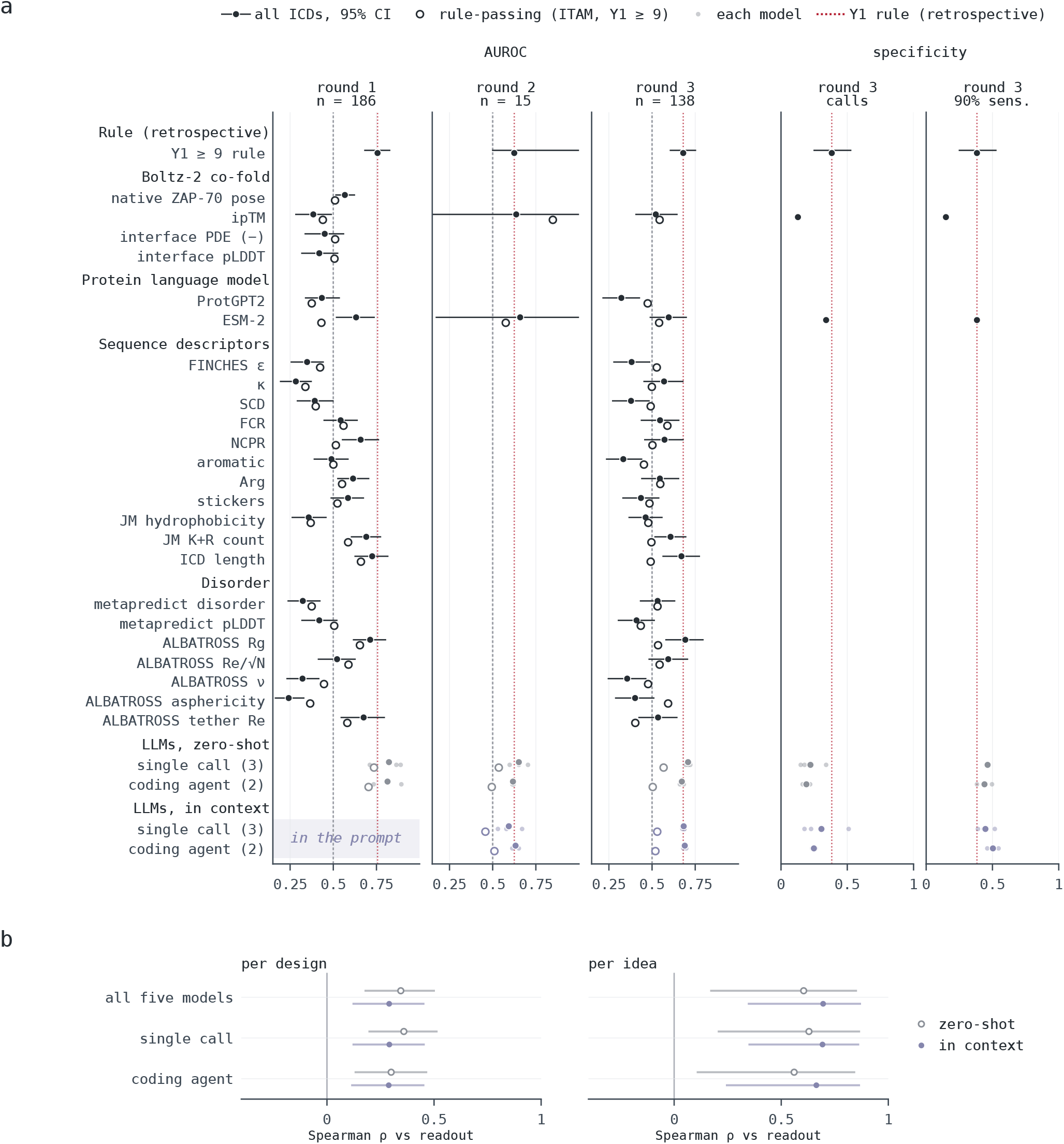
Benchmarks of the model predictions. **a**, Every method on the same ICDs: the 186 library ICDs of round 1, the 15 new designs of round 2 and the 138 new ICDs of round 3. In round 2, only the Y1 rule, Boltz-2 ipTM, ESM-2 and the language models were scored. AUROC for active versus inactive ICDs, for all ICDs (filled, with 95% bootstrap interval) and for ICDs that pass the Y1 rule (open), and round 3 specificity of each method’s own calls and at the threshold that keeps 90% of active ICDs (90% sens.), shown for the methods that give calls or a stored score. Scores include the Y1 rule, Boltz-2 co-fold scores, protein language-model likelihoods, and condensate, charge, composition and disorder descriptors. Language models, the mean over the single-call LLMs (GPT-6.1, Gemini 4, Claude Opus 5) or the coding agents (Claude Code, Codex), with each model as a small dot; in context, the prompt held the labeled ICDs of rounds 1 and 2 (shaded, round 1 in the prompt). The Y1 rule was derived retrospectively, after all screens were measured, and is shown as a reference rather than a prediction. Dashed line, chance; dotted line, the Y1 rule. **b**, Spearman correlation of P(active) with the activity readout (round 2, IFN-γ relative to `BBζ` (`4-1BB–CD3ζ`); round 3, killing relative to `krgr–ζ1–40`), per design and per idea, zero-shot (open) and in context (filled), for all five models, the single-call LLMs and the coding agents; bars, 95% bootstrap intervals.

**Figure S9.**
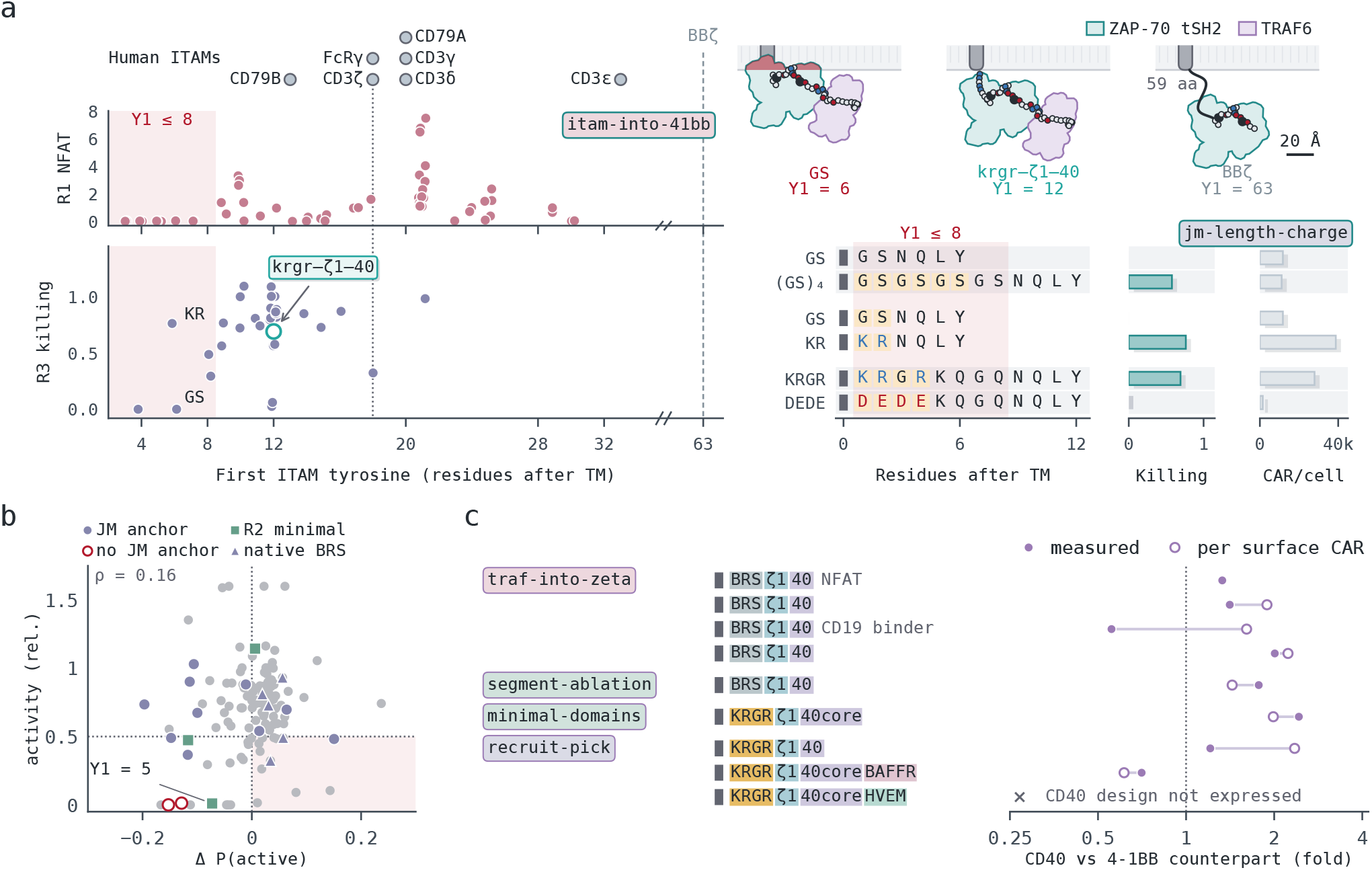
Membrane proximity, model predictions and the CD40 core. **a**, Membrane proximity: Y1 of human ITAM-bearing chains, round 1 NFAT and round 3 killing; shading, Y1 of 8 or less. Right, JM contrasts with killing and surface CAR/cell. Top right, Boltz-2 models of ZAP-70 and TRAF6 docked at Y1 = 6, 12 and 63. **b**, Whether the models’ update tracked the outcome. Change in the ensemble’s P(active) from zero-shot to in context (the five models as one ensemble, as in Fig. 5c) against the activity readout relative to the reference (round 2, IFN-γ; round 3, killing relative to `krgr–ζ1–40`), for the 125 designs scored in both tasks (gray), with the cases of Fig. 5b on top: shortened round 3 designs with (filled circles) or without (open circles) a JM segment before the first module, round 2 minimal domains (squares) and round 3 native-BRS JM designs (triangles). Dotted lines, zero change and the 0.5 activity threshold; shading, raised in context but inactive. Values above 1.6 are drawn at 1.6. **c**, The CD40-core thread: each CD40 design relative to its 4-1BB counterpart, measured (filled) and per surface CAR (open). One measurement per round 3 construct.

#### Settings

All five models received one task text (T1). It describes the CAR, the three assays and the files: test.csv, the ICDs to predict, and in context train.csv, the labeled ICDs of the earlier rounds. The single-call LLMs received the same text with the files inline and answered in a JSON schema; the coding agents worked in an isolated sandbox that held the files and wrote predictions.csv (T2). Each prediction comprises a short reasoning, a probability and a label, and each model was run three times per condition. Zero-shot, a model saw no labels; in context, it saw the labeled ICDs of the earlier rounds (round 1, 186 ICDs, for the 15 round 2 designs; rounds 1 and 2, 279 labeled measurements of 201 unique ICDs, for the 138 round 3 ICDs). Every run returned a prediction for every ICD.

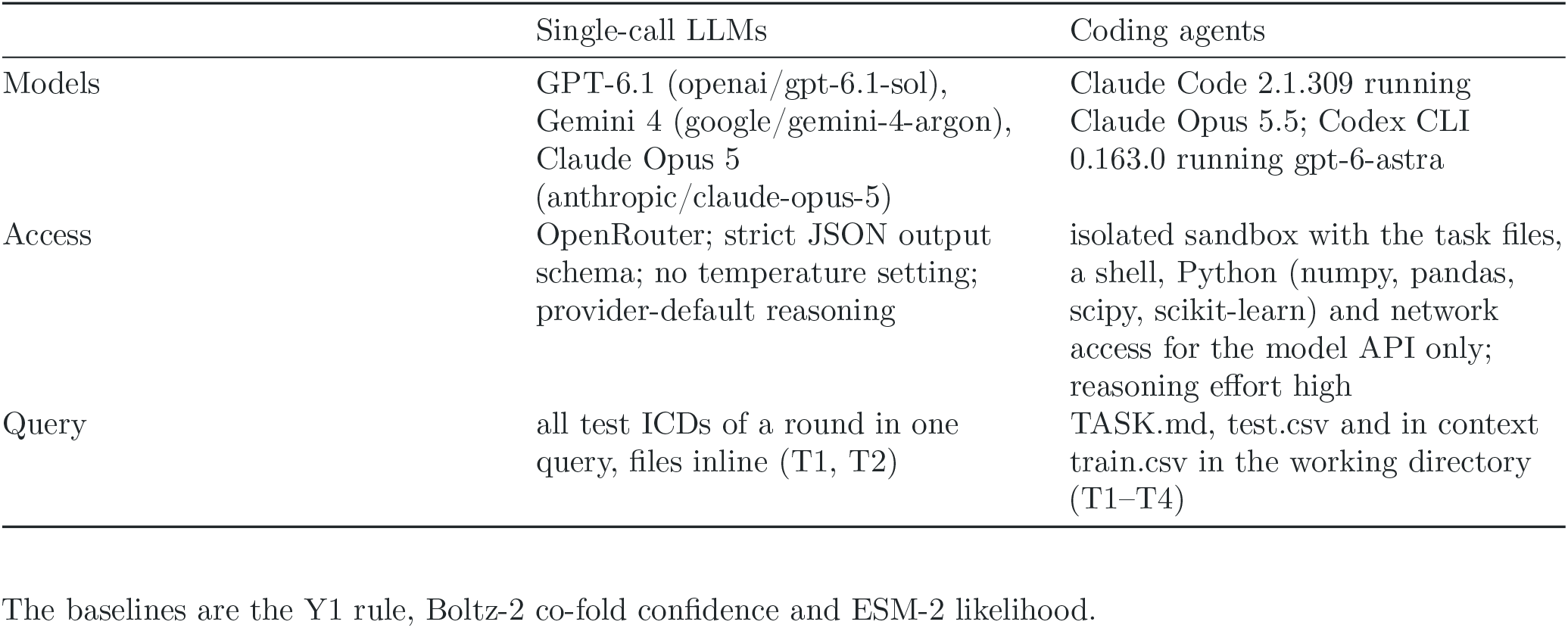

The baselines are the Y1 rule, Boltz-2 co-fold confidence and ESM-2 likelihood.

#### Comparator task

In a second task (C1), the same five models estimated the probability that each ICD’s readout exceeded that of the round’s comparator on the same plate, P(better): `BBζ` for the 15 round 2 designs and `krgr–ζ1–40` for the 117 round 3 designs with an IFN-γ measurement. The task names the comparator and gives its sequence; the round 3 readout is IFN-γ, as in the round 3 screen. In context, train.csv gave each earlier measurement’s round, assay, ratio to `BBζ` and label (higher, a ratio of 1 or more, or lower): the round 1 library in Jurkat NFAT cells and the 78 library designs re-screened in primary T cells (264 measurements) for round 2, with the round 2 plate added (282 measurements) for round 3. The single-call LLMs ran with reasoning effort high, Claude Opus 5 through the Claude API with adaptive thinking; each model ran three times per condition.

#### Scoring

AUROC and specificity were computed per run against the measured calls and averaged over runs, with 95% intervals from 1,000 bootstrap resamples of ICDs. Specificity was computed for each model’s own calls and at the highest threshold on its predicted probability that still called at least 90% of active ICDs active; the latter, like AUROC, depends only on the order of the predicted probabilities, not on where the model places its call threshold.

#### Prompts

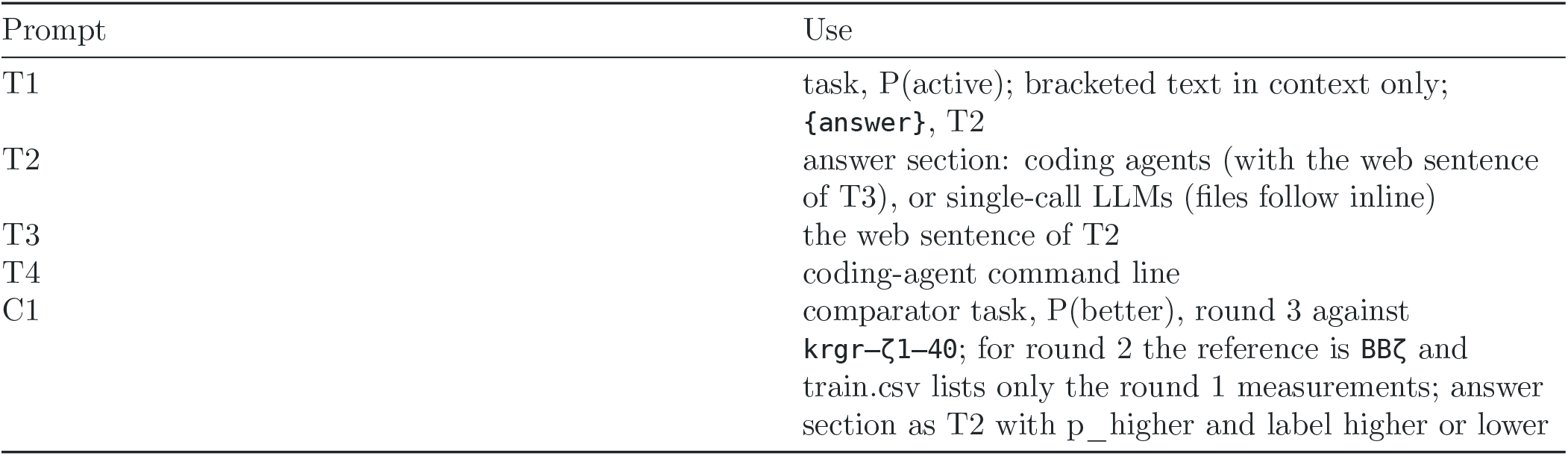

**T1**

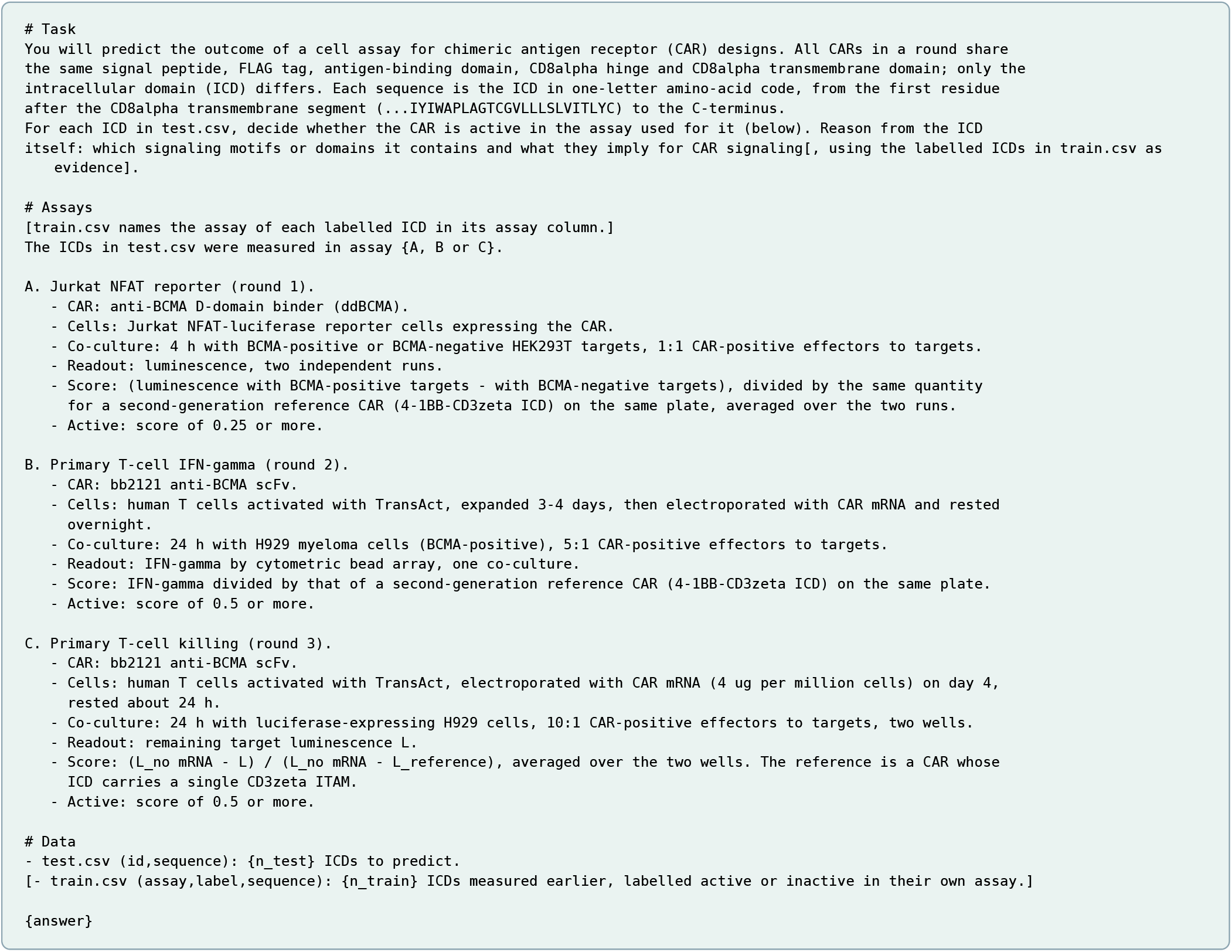

**T2**

**Coding agents**

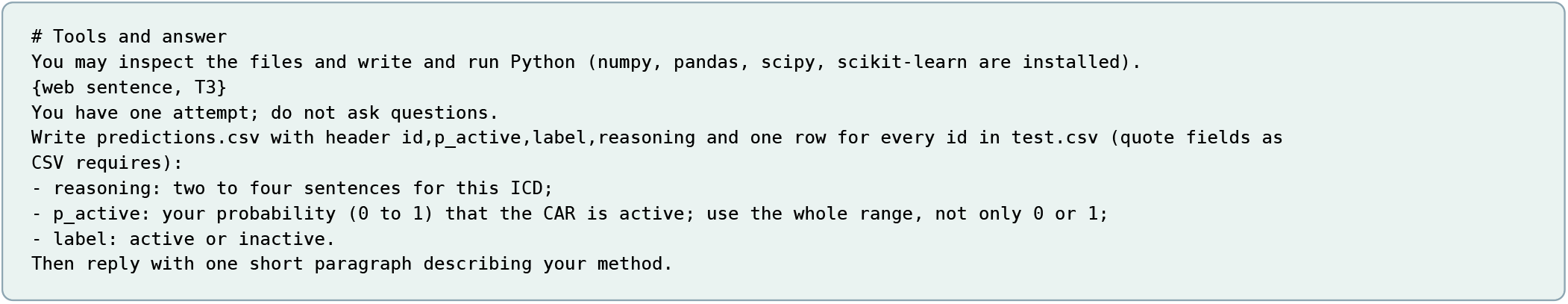

**Single-call LLMs**

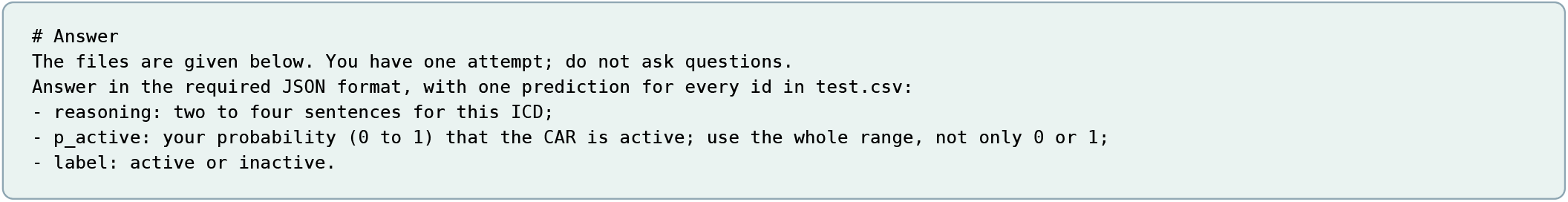

**T3**

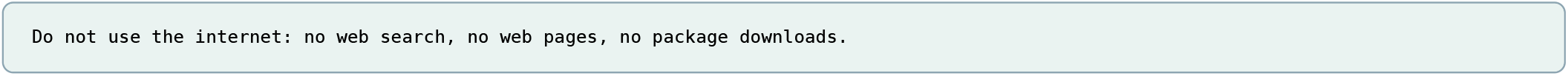

**T4**

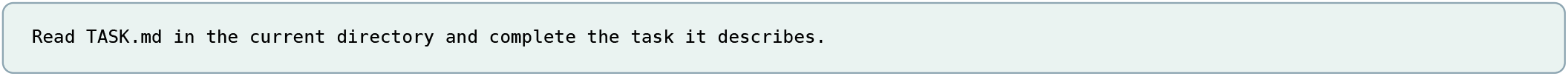

**C1**

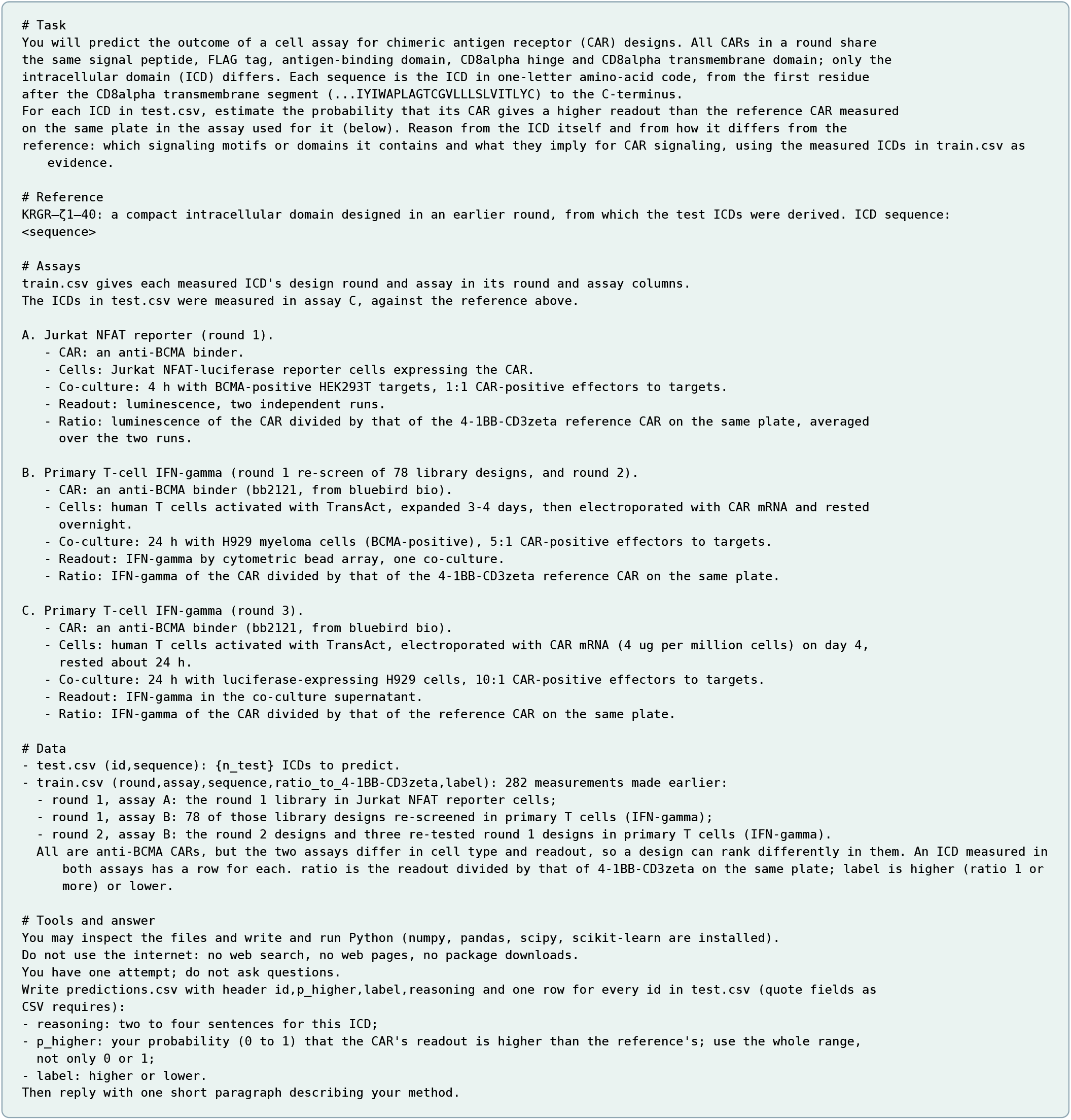

**Figure S10.**
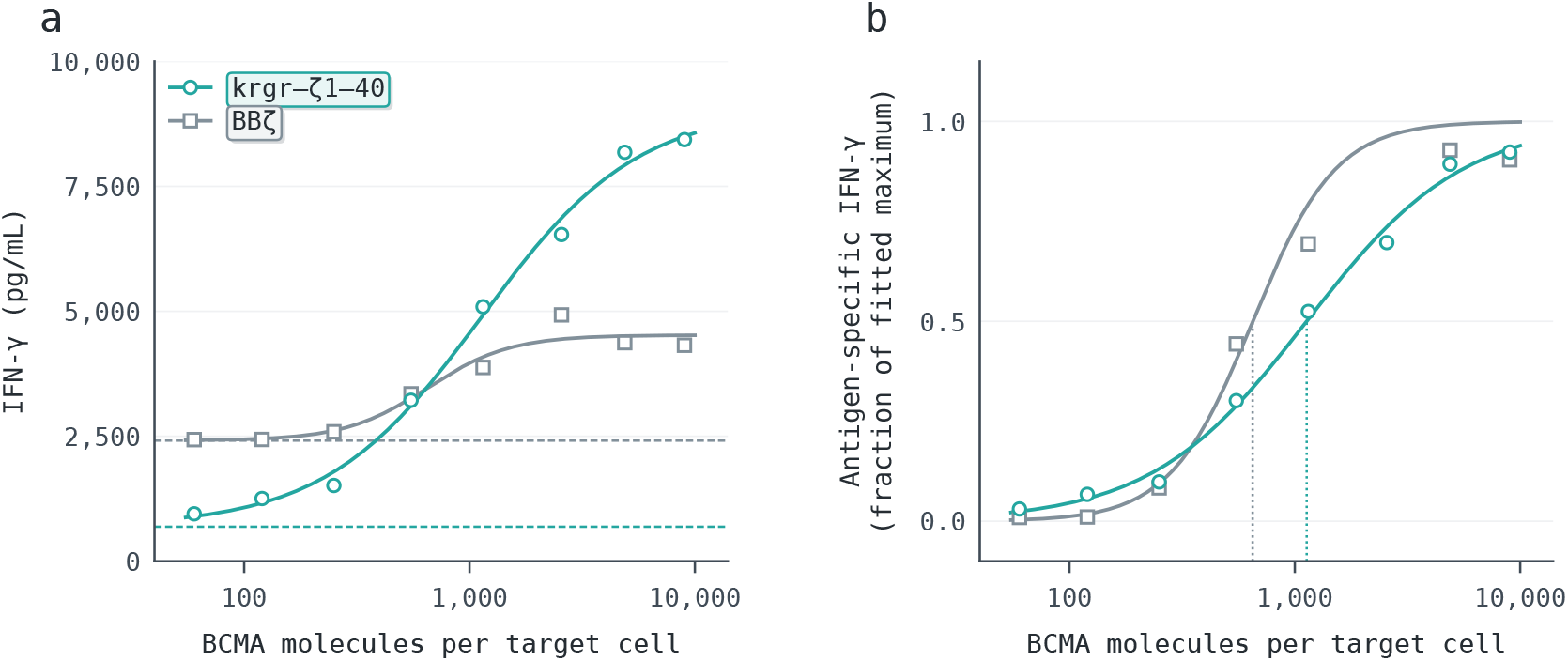
Antigen sensitivity of IFN-γ release at a matched CAR mRNA dose. **a**, IFN-γ from `BBζ` (`4-1BB– CD3ζ`) and `krgr–ζ1–40` CAR T cells after co-culture with target cells carrying eight densities of BCMA, at the highest mRNA dose of Fig. 4a; dashed lines, each construct’s fitted antigen-independent floor. **b**, The same readings with the fitted floor subtracted, as a fraction of each fitted span; dotted lines, EC50s of 650 (`BBζ`) and 1,130 (`krgr–ζ1–40`) BCMA molecules per target cell.

### Structural and sequence analyses of CAR ICDs

To examine the structural context of ICD activity, we modeled adaptor binding and membrane interactions. Predicted complexes reproduced native ZAP-70–ITAM and TRAF6–CD40 binding poses for active and inactive ICDs alike, so binding geometry did not explain activity (Fig. S11a–e).

**Figure S11.**
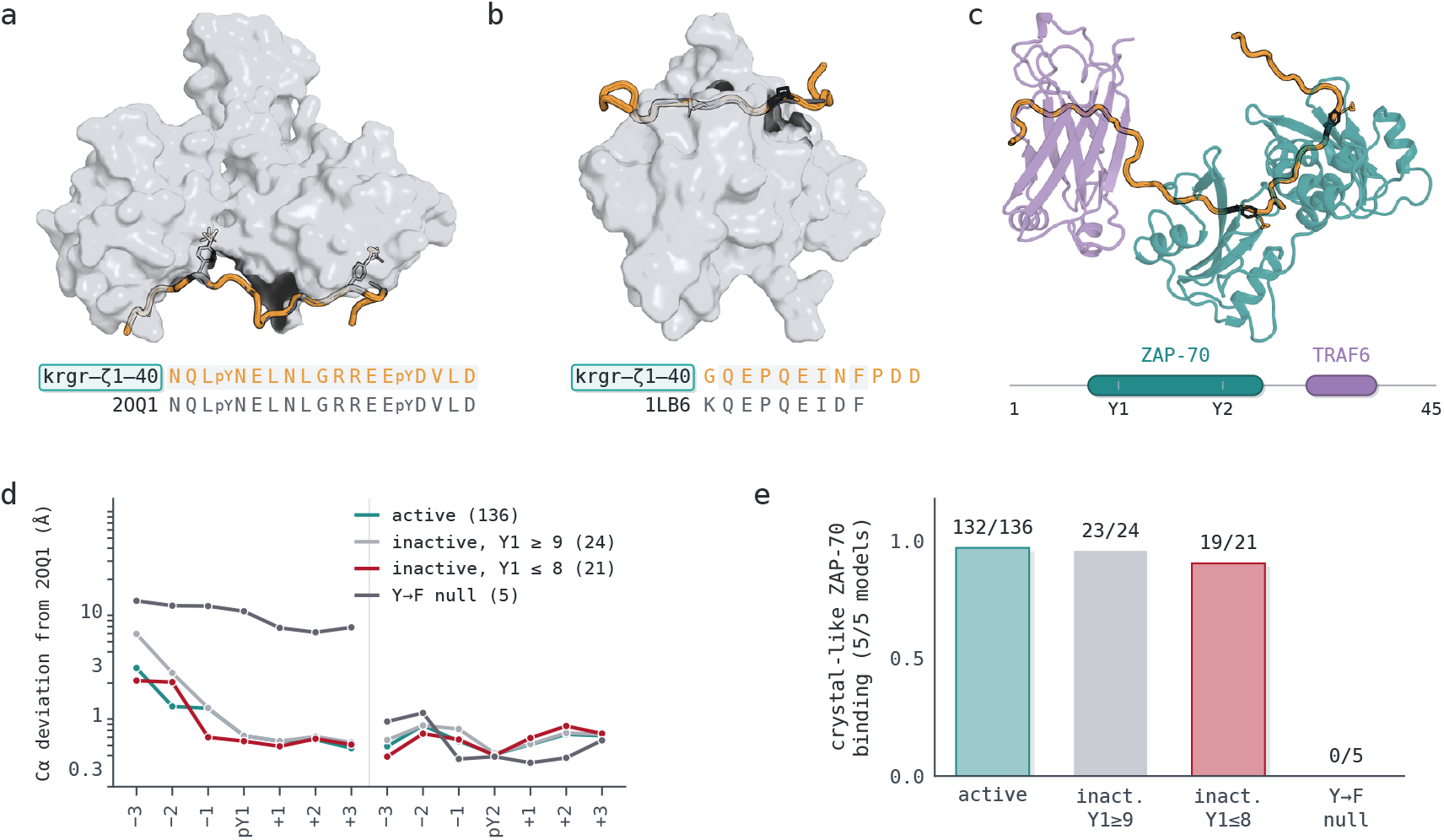
Structure models of krgr–ζ1–40 and related ICDs. **a**, Boltz-2 co-fold of `krgr–ζ1–40` with the ZAP-70 tandem SH2 domains, superposed on PDB 2OQ1. **b**, Co-fold of the CD40-derived segment with the TRAF6 TRAF-C domain, superposed on PDB 1LB6. **c**, Ternary co-fold of the ICD with ZAP-70 (teal) and TRAF6 (mauve), which bind separate segments. **d**, Deviation of each ITAM half-site from 2OQ1 for active, inactive and Y→F null ICDs; only the nulls are displaced. **e**, Fraction of library ICDs whose first ITAM is crystal-like in all five models, by group.

We next asked whether natural sequences and physical models could explain the spacing constraint. Membrane-proximal tyrosines in human single-pass proteins were less frequently annotated or detected as phosphorylated (Fig. S12a–c). Lck substrate scores (Fig. S12f) and a steric model of ZAP-70 access (Fig. S12k) did not account for the sharp activity boundary observed in round 1.

**Figure S12.**
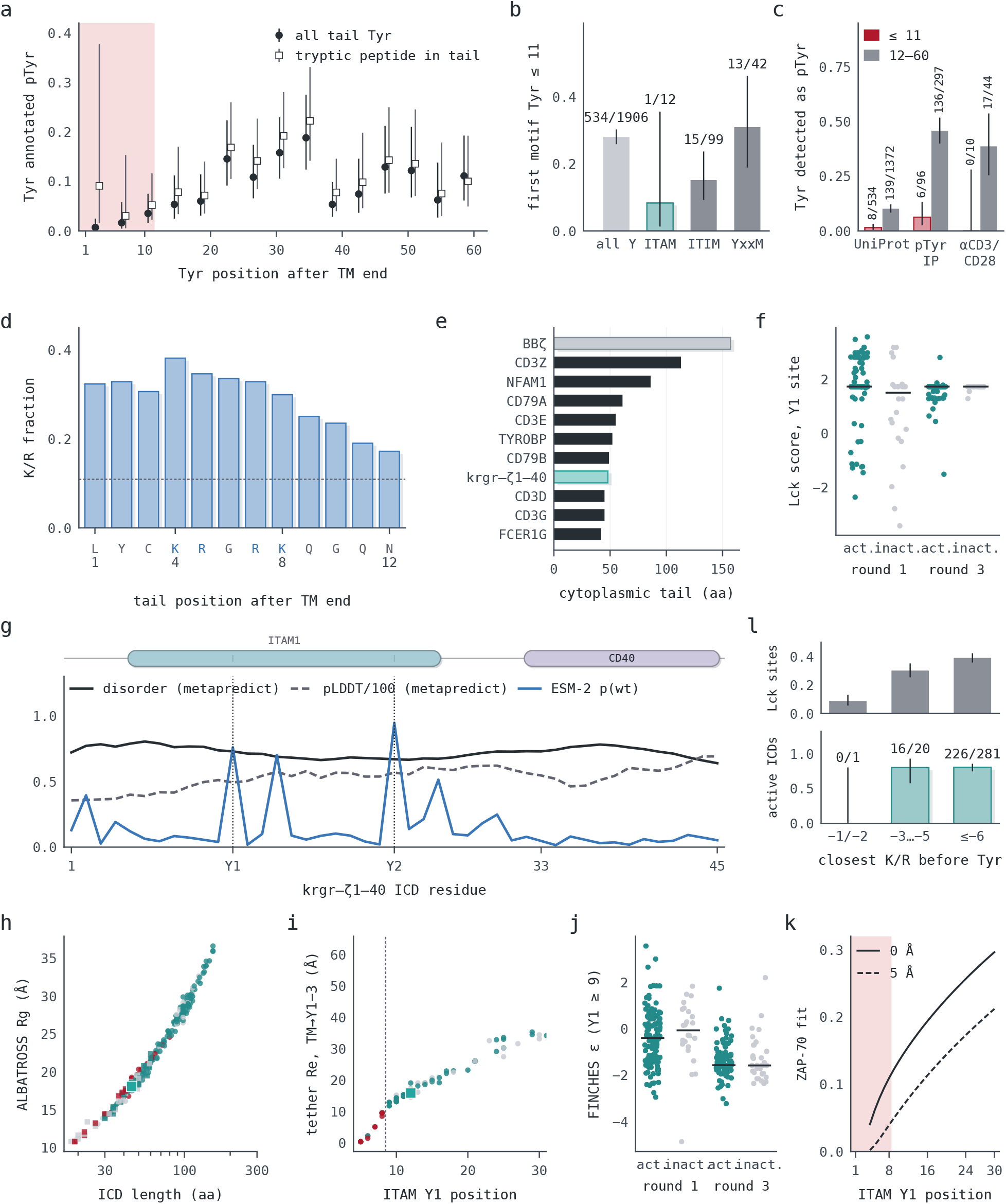
Sequence, proteome and physical-model context for the juxtamembrane spacing. **a**, Fraction of cytoplasmic tyrosines annotated as phosphotyrosine in human single-pass proteins by distance from the transmembrane end, for all tail tyrosines (filled circles) and those whose tryptic peptide lies wholly in the tail (open squares). **b**, Fraction of motif tyrosines within 11 residues of the membrane for all tyrosines, ITAMs, ITIMs and YxxM motifs. **c**, Phosphotyrosine detection within 11 residues (crimson) and at 12–60 (gray) in UniProt and in two Jurkat phosphotyrosine datasets [132,133]. **d**, K/R fraction at each tail position. **e**, Cytoplasmic tail lengths of natural ITAM chains, `krgr–ζ1–40` and `BBζ` (`4-1BB–CD3ζ`). **f**, Lck substrate score of the Y1 site [134] for active and inactive ICDs with Y1 of 9 or more. **g**, Per-residue disorder, predicted pLDDT and ESM-2 wild-type probability along `krgr–ζ1–40`. **h**, Radius of gyration against ICD length (teal, active; crimson, Y1 of 8 or less; gray, other inactive; square, `krgr–ζ1–40`). **i**, End-to-end distance from residue 1 to Y1 − 3 against Y1. **j**, Homotypic interaction strength (FINCHES ε [135]) for ICDs with Y1 of 9 or more. **k**, Probability that the 2OQ1 complex fits below the membrane in a steric model, against Y1; the model rises gradually rather than stepping at Y1 = 9 as in Fig. S9a. **l**, Efficient Lck phosphorylation of library substrate sites (top) and the fraction of active ICDs (bottom) by the position of the nearest K/R before the tyrosine [136]. Positions in a–e count from the UniProt transmembrane end, three residues earlier than in Fig. S9a.

### Agentic digital infrastructure

We built an electronic lab notebook system to store all our sequences, protocols, and experimental data. Claude Code serves as the primary interface, with dedicated graphical interfaces for tasks that demand consistent visual interaction, such as plate map visualization and flow cytometry gating.

Routine procedures were written as shared agent skills instead of fixed application code. For example, coding agents performed codon optimization from a shared skill that defined the procedure and adapted it to individual sequences and constraints, without each edge case being encoded in advance [137,138].

We use minimal data models to store only core information and leverage agents to infer many relationships and details at runtime (Fig. S13). For example, after each experiment, users upload raw data to a shared filesystem, where agents use plate maps to link measurements to designed variants and experimental conditions for analysis and plotting. Typically, we discuss results and make decisions with comprehensive analyses and plots in hand within minutes of instrument readout.

The PostgreSQL database and file storage ran on Supabase.

**Figure S13.**
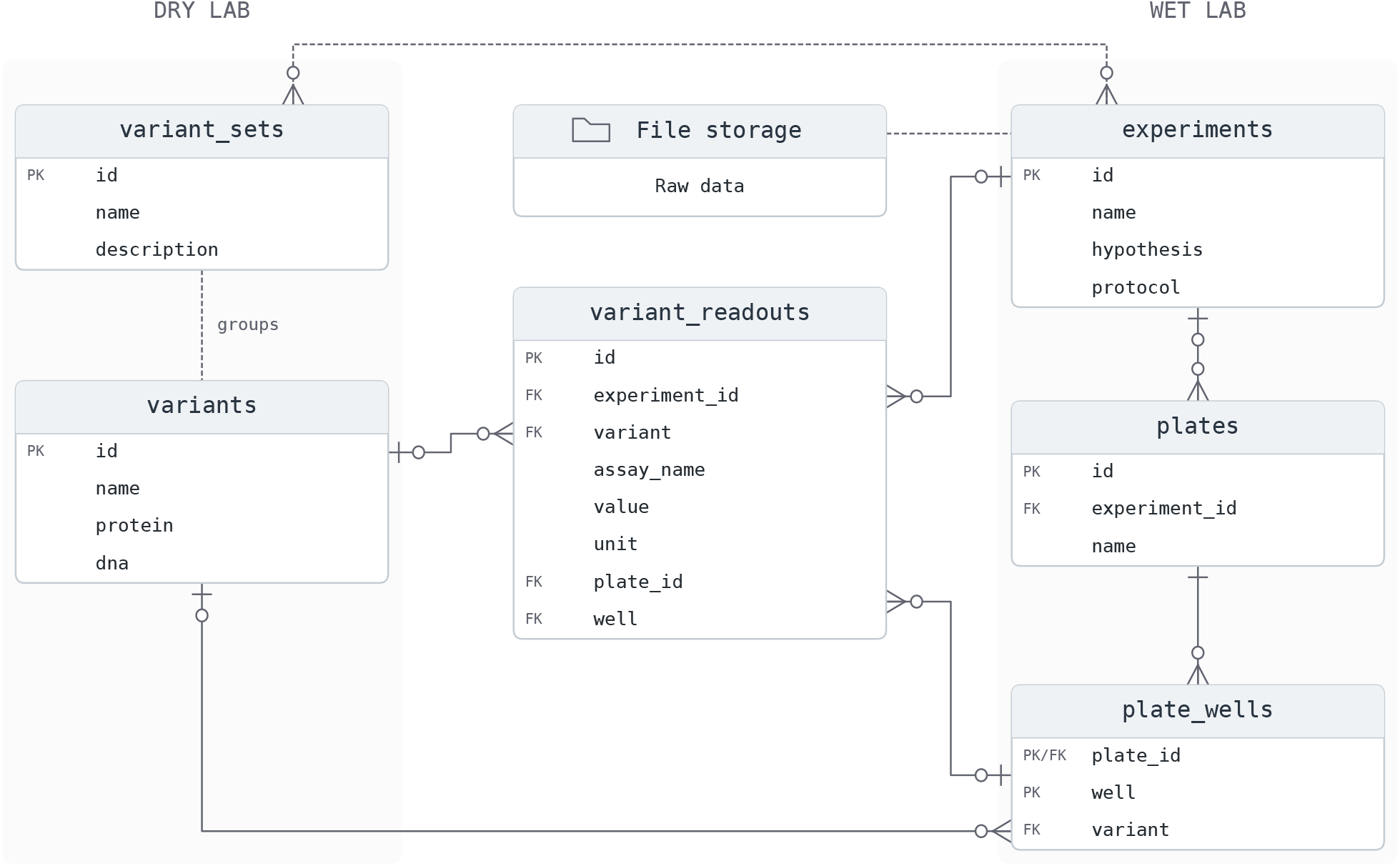
Electronic lab notebook data model. Selected tables and fields link experiments, variants, plates, and readouts. Computational scientists typically populate `variants` after sequence design; wet-lab scientists create `experiments` and `plates` during experiment planning. Agents read and write all tables. `variant_readouts`, populated by agents, links variants to experimental measurements for analysis. PK, primary key; FK, foreign key.

**Supplementary Table 1.**
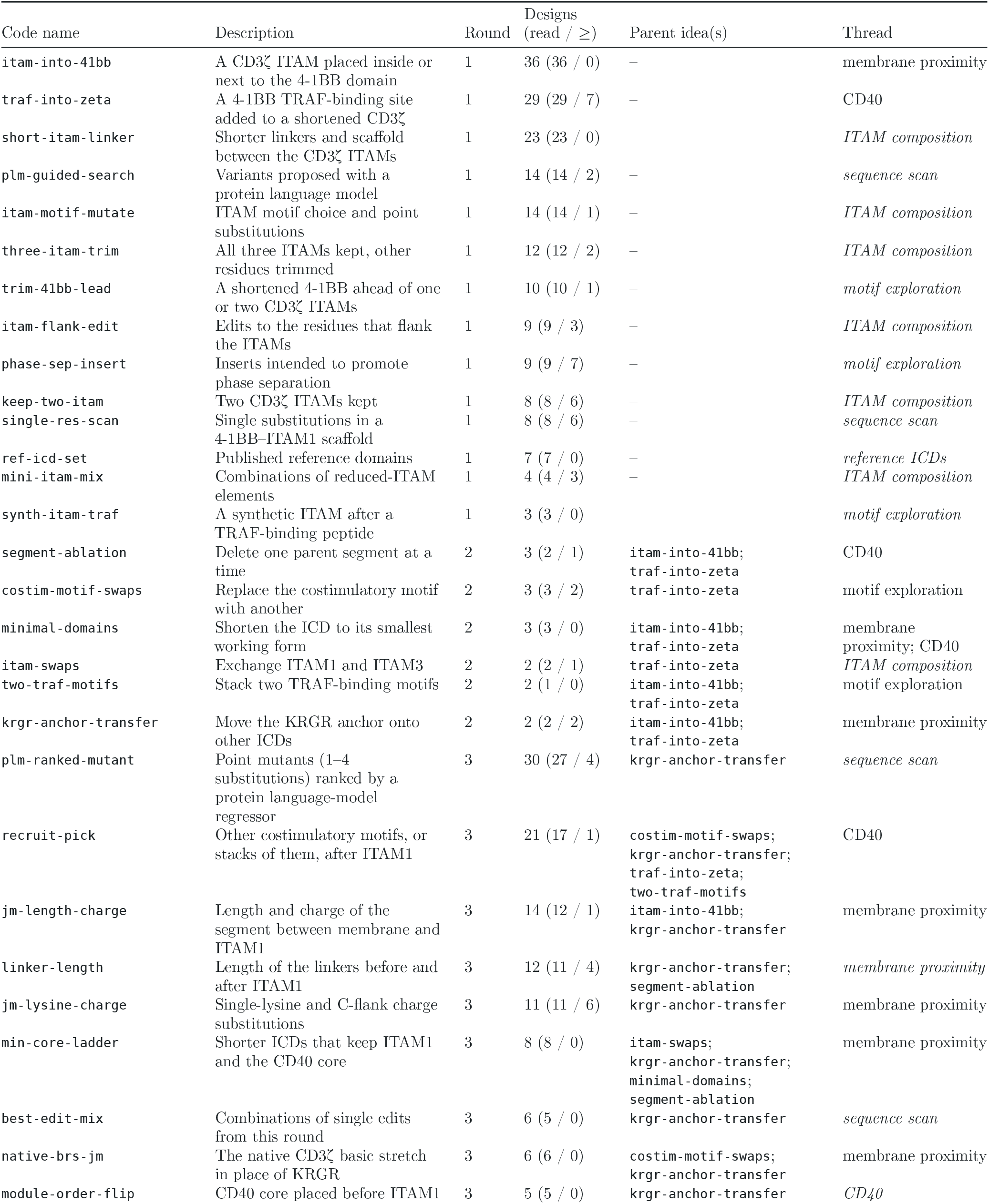
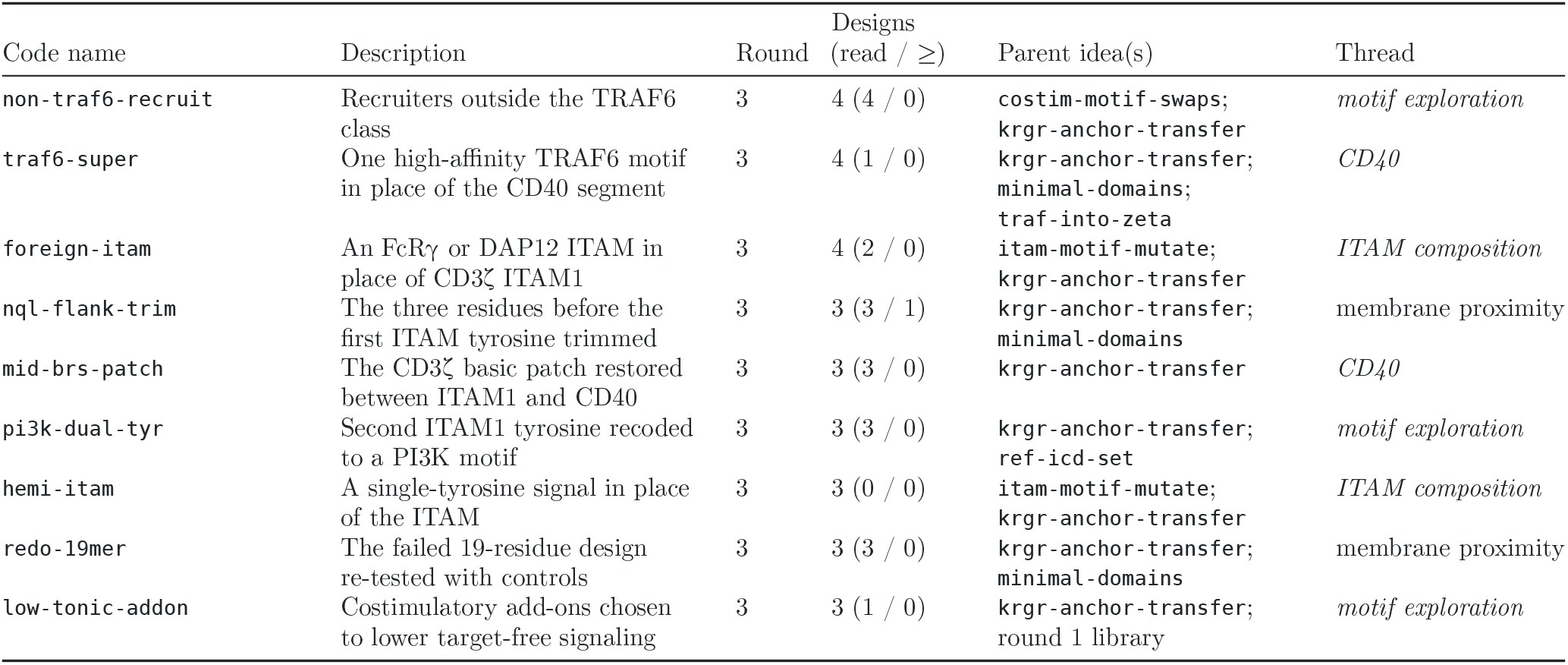
The 38 ideas of Fig. 5a. The ideas, their code names and descriptions, and this table were generated by AI agents. The full table (CSV; Markdown) gives, for each idea, its code name, round, plain-language description, short recorded intention, designs, designs with a readout and designs at or above the comparator, its parent ideas with the evidence type of each link, and the Fig. 5a thread it is on (ITAM distance from membrane, teal; CD40 core, lavender; both; or none). The short version below gives each code name with a brief description and its round. Designs are listed with, in parentheses, those with a readout and those at or above the comparator; comparators are as in Fig. 5a. Parent ideas are linked by a parent design, a KRGR-anchor donor, a citation in the design rationale or an inferred shared edit, and every round 3 idea also descends from `krgr–ζ1–40` in `krgr-anchor-transfer`. Threads are named as in the Fig. 5a key; ideas off the drawn threads are assigned from their description and shown in italics (ITAM composition, motif exploration, sequence scan and reference ICDs are themes, not Fig. 5a threads).

### Round 1 design prompts

We recovered 17 distinct prompts from 34 archived round 1 agent runs.

#### Model assignments

Selected model assignments are listed below.

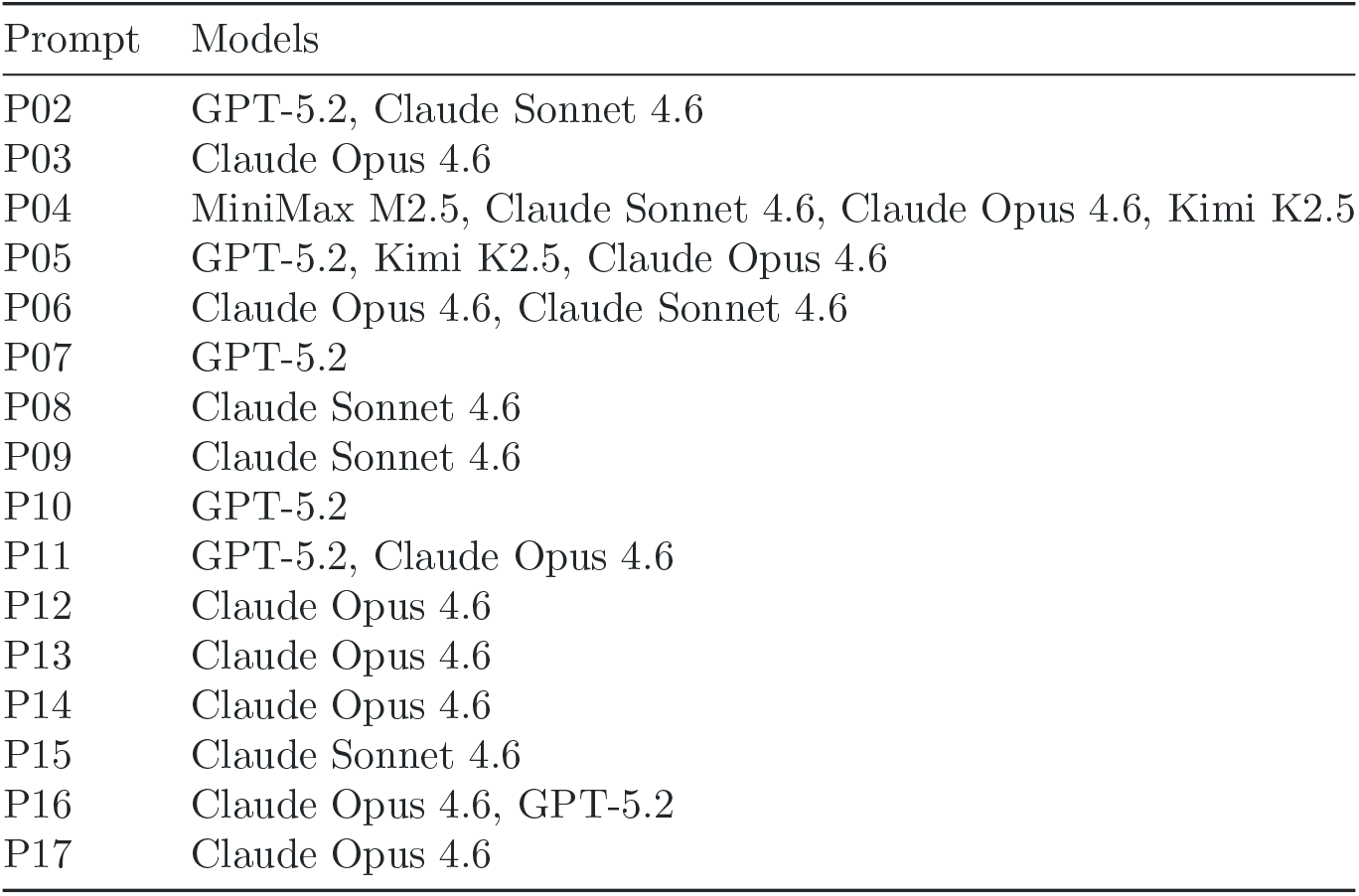

#### Prompts

The task-specific text below was embedded in the system message. Each run also received the following instruction, followed by an automatically generated execution plan:

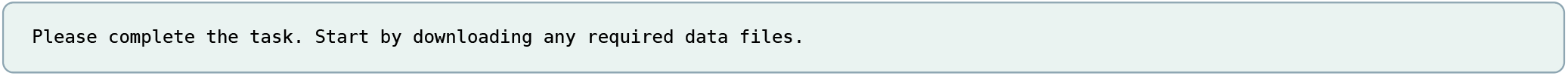

**P01**

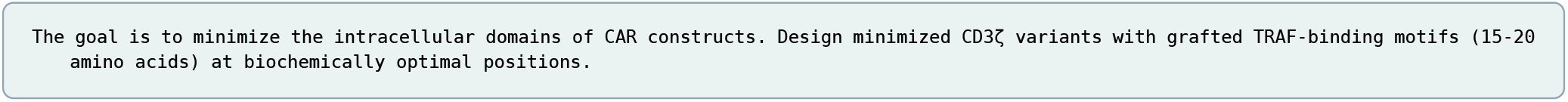

**P02**

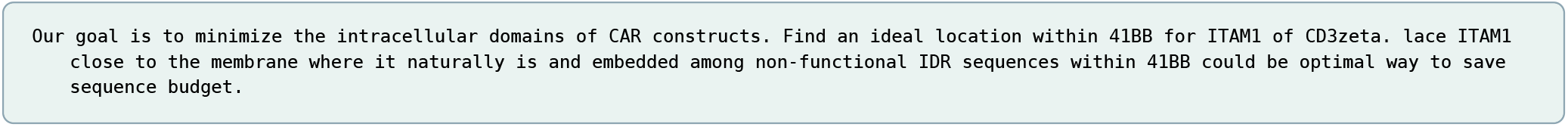

**P03**

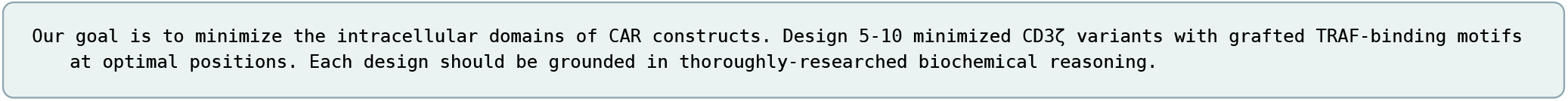

**P04**

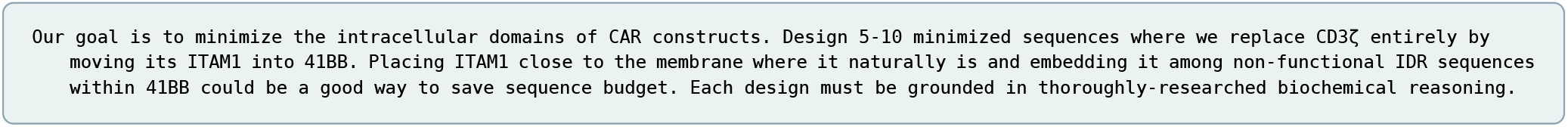

**P05**

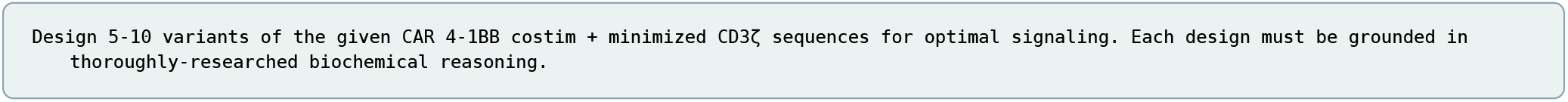

**P06**

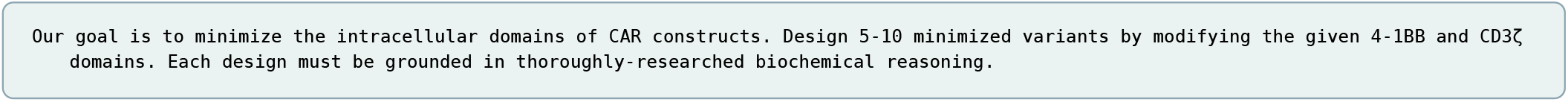

**P07**

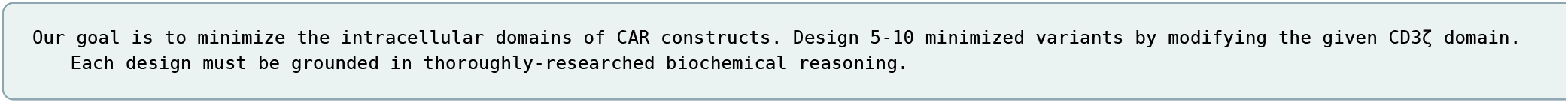

**P08**

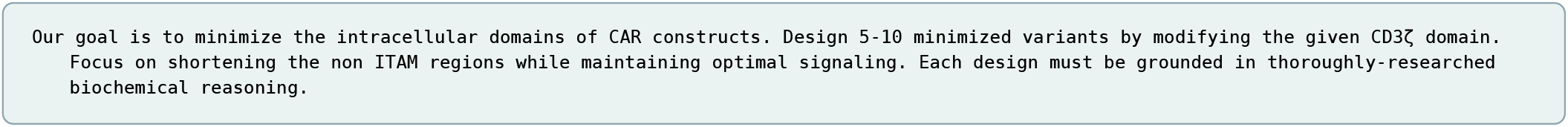

**P09**

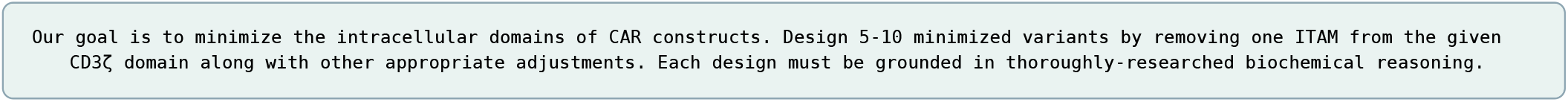

**P10**

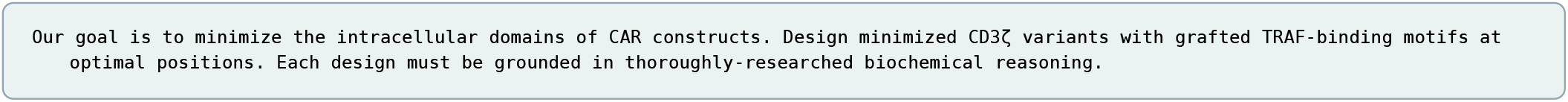

**P11**

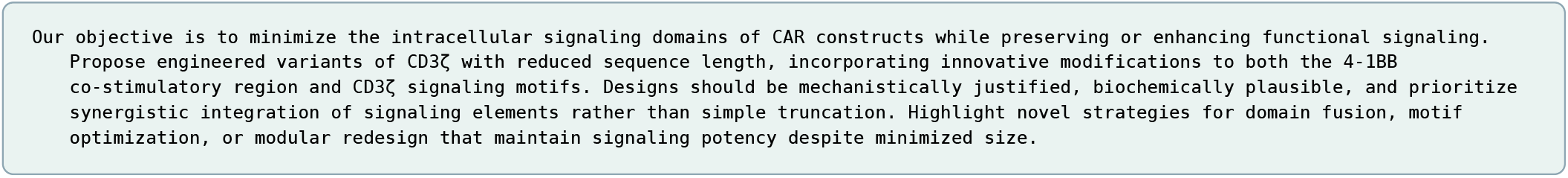

**P12**

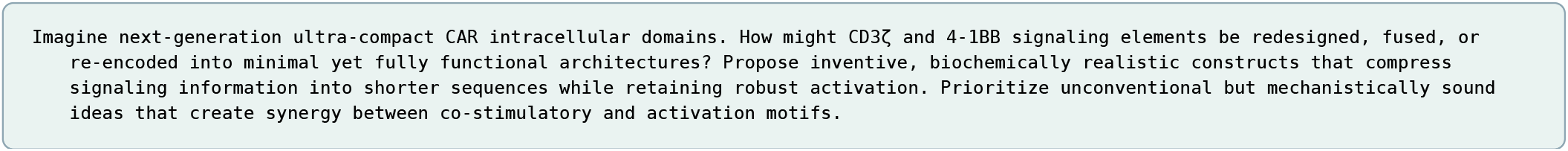

**P13**

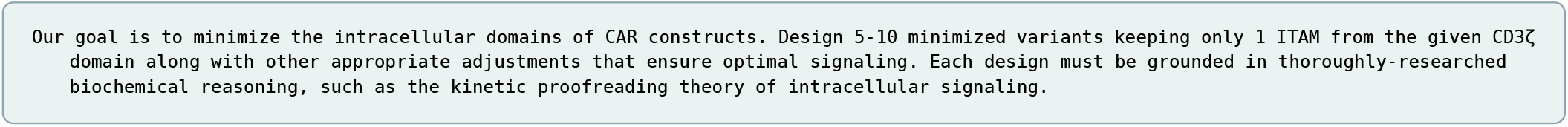

**P14**

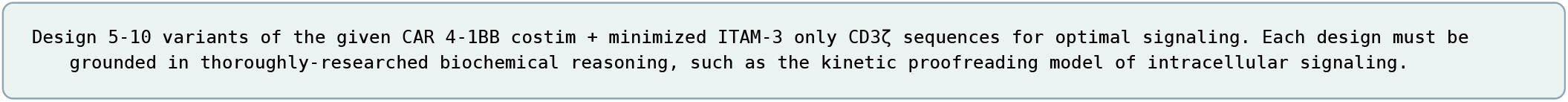

**P15**

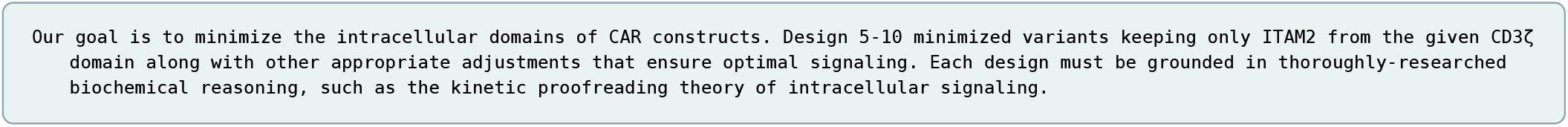

**P16**

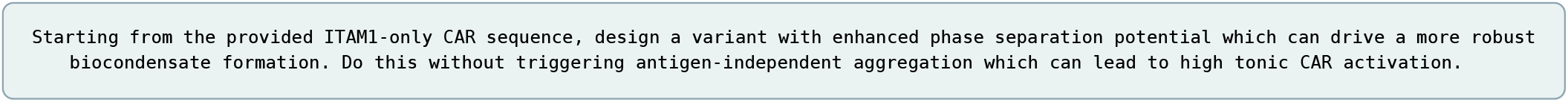

**P17**

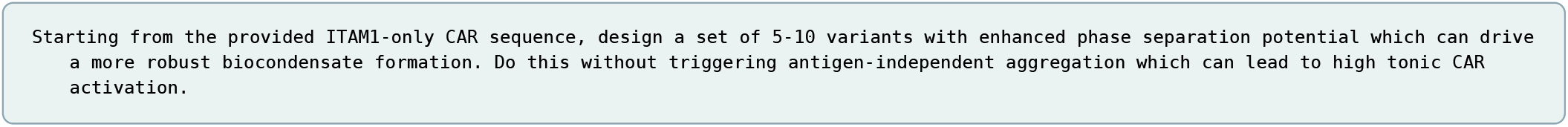

